# Scale-Aware Compositional Inference Improves Reproducibility and Uncovers Convergent Aging Programs in Spatial Transcriptomics

**DOI:** 10.64898/2026.07.27.740958

**Authors:** Deniz Parmaksiz, Steffy B. Manjila, Kyle C. McGovern, Donghui Shin, Ingvild E. Bjerke, Anirban Paul, Justin Silverman, Yongsoo Kim

**Affiliations:** Department of Neuroscience and Experimental Therapeutics, College of Medicine, The Pennsylvania State University, Hershey, PA, USA; Neuroscience Center of Excellence, School of Medicine, Louisiana State University Health Sciences Center, New Orleans, LA, USA; College of Information Sciences and Technology, The Pennsylvania State University, University Park, PA, USA, 16802; Department of Biological Sciences, College of Science, Purdue University, West Lafayette, IN, USA, 47906; Program in Bioinformatics and Genomics, The Pennsylvania State University, University Park, PA, USA, 16802

**Keywords:** Spatial transcriptomics, Differential expression, Aging, MERFISH, Compositional data analysis, Cross-study reproducibility

## Abstract

Spatial transcriptomics enables analysis of molecular organization with anatomical context. Existing spatial differential expression methods are restricted to within-sample inference, forcing between-sample comparisons to rely on approaches adapted from single-cell RNA-seq. Here, we establish a scale-aware inference framework for spatial differential expression by modeling compositional constraints and variation in total RNA abundance rather than removing them through normalization, enabling calibrated between-sample inference at cell-level resolution. Our method produces more reliable results in simulated data and different spatial platforms. When applied to aged mouse brains, the analysis reveals converging aging-associated programs involving cellular signaling, membrane homeostasis, and neurovasculature across independent datasets.

## Background

Spatial transcriptomics (ST) technologies enable molecular profiling while preserving spatial context within intact tissue, providing a powerful framework for studying tissue organization and molecular programs across biological and pathological conditions (Park et al., 2023; Piwecka et al., 2023). Differential-expression (DE) analysis is one of the primary statistical tools used to compare biological groups or conditions in ST. Broadly, these analyses fall into two settings: studies in which the variable of interest varies within a tissue section (e.g., anatomical regions or cellular neighborhoods) and replicated studies in which biological conditions such as age, disease, genotype, or treatment vary between animals or tissue sections. The latter presents a distinct inferential problem because the condition of interest varies between biological samples, while thousands of individual cells are observed within each sample. Treating these cells as independent experimental units can substantially overstate the available information. Pseudobulk analysis restores the biological sample as the unit of inference but discards cell-level heterogeneity and spatially resolved annotations (Squair et al., 2021; Tung et al., 2017). Replicated ST studies therefore require methods that respect sample-level replication without unnecessarily collapsing cell-resolved measurements.

Accounting for repeated measurements addresses the unit-of-inference problem, but it does not resolve a separate challenge arising from count-based expression data: observed counts alone cannot determine whether an apparent gene-specific change reflects true regulation or a broader shift in overall transcript abundance (referred to here as transcriptional scale). This challenge is particularly relevant when studying biological conditions that produce coordinated transcriptome-wide changes. Aging, for example, causes broad reductions in total transcriptional output alongside gene-specific changes (Stoeger et al., 2022). If one set of genes is downregulated while another remains unchanged, the unchanged genes will falsely appear upregulated despite no change in their absolute expression. Conversely, coordinated changes across many genes may be difficult to detect if their relative proportions remain similar. In imaging-based ST, this scale uncertainty is further compounded because the total detected transcripts reflects inseparable biological and technical factors, including panel size and composition, detection efficiency, sample quality, assay chemistry, and biological state (Bhuva et al., 2024; Grases & Porta-Pardo, 2026).

Most DE workflows address scale ambiguity through normalization. Library totals or estimated size factors are used to place cells or samples onto a common scale, after which normalized values are analyzed as though they were directly comparable. Although often described as removing technical variation, normalization cannot recover unobserved biological transcriptomic scale from count data alone. Instead, it resolves the ambiguity by imposing fixed assumptions about how total transcript abundance varies across cells or conditions. Normalization-based workflows therefore treat these estimated scaling factors as known rather than propagating uncertainty about them into downstream inference (Nixon et al., 2025; McGovern & Silverman, 2026). When these normalization assumptions are incorrect, broader transcriptome-wide changes may be misinterpreted as gene-specific regulation, while true coordinated biological shifts may be attenuated or removed entirely.

In imaging-based targeted assays, count-based normalization can additionally induce region- and cell-type-specific bias in DE statistics and effect-size estimates when targeted panel composition constrains the set of observable genes (Atta et al., 2024). Recent efforts have sought to address these challenges by improving normalization rather than DE inference itself. Spatially varying normalization methods, such as SpaNorm, adapt preprocessing to account for spatially structured library-size effects (C. Li et al., 2023; Salim et al., 2022; Wang et al., 2022; Salim et al., 2025). However, these methods still produce a single deterministic normalized expression matrix and therefore do not explicitly propagate uncertainty into downstream DE modeling.

While normalization methods focus on preprocessing, other ST-specific methods instead model spatial dependence, addressing a challenge that is distinct from transcriptomic scale uncertainty. Frameworks such as STdiff and SpatialGEE model residual dependence among nearby cells or spots, while smiDE accounts for transcript misassignment resulting from imperfect cellular segmentation (Ospina et al., 2024; Vasconcelos et al., 2026; Wang et al., 2026). Other approaches target cell-type-specific expression in multicellular spots or expression associated with defined cellular neighborhoods (Cable et al., 2022; Mason et al., 2024). These methods are particularly useful when the predictor of interest varies within a tissue section or when local spatial dependence is the primary focus. They do not, however, resolve transcriptomic scale uncertainty or account for the repeated-measures structure of multi-sample studies (e.g., comparing age, treatment, or disease state across animals). Spatial dependence, biological replication, and unmeasured scale represent three distinct, though potentially interacting, sources of uncertainty.

As ST datasets rapidly accumulate across platforms, laboratories, and experimental conditions, the scale assumptions underlying DE analyses have emerged as a critical barrier to cross-study reproducibility (Zhou et al., 2023; Plummer et al., 2025). Differences in assay chemistry, panel composition, tissue sampling, and detection efficiency fundamentally alter the relationship between observed counts and underlying transcript abundance. Consequently, standard normalization strategies may resolve transcriptomic scale differently across datasets, producing discordant gene-level conclusions even when the underlying biology is similar. Cross-study reproducibility thus serves as a practical test of whether a DE framework recovers biological signals that remain interpretable and consistent across distinct experimental settings.

Here, we establish scale-aware inference framework for replicated, cell-level ST studies based on the ANOVA-Like Differential Expression (ALDEx) 3 mixed-effect model (McGovern and Silverman, 2026). The framework propagates uncertainty in both relative transcript composition and global transcript abundance, while mixed-effects models account for the correlation of cells within biological samples and imaging runs. Using simulations together with independent MERFISH and CosMx mouse-brain aging datasets, we evaluate statistical calibration, cross-study reproducibility, and biological concordance. We identify reproducible glial and vascular programs involving cellular transport, extracellular matrix remodeling, and trophic signaling, demonstrating a significant advantage of scale-aware count-compositional inference across imaging-based ST technologies.

## Results

### Spatial transcriptomics data is count compositional

Targeted single-cell ST assays generate data that are fundamentally count-compositional: observed counts are measured from a predefined set of genes, embedded within spatial tissue architecture, and reflect uncertainty in both the total number of detected transcripts per cell and the relative composition of transcripts across neighboring cells.

#### Scale uncertainty

The first component of count-compositional uncertainty concerns global scale, representing the total transcript abundance underlying the observed counts. Because imaging-based ST measures only a targeted subset of the transcriptome, observed transcript totals cannot generally be interpreted as direct measurements of absolute RNA abundance. DE analyses relying on conventional normalization therefore assume that a single scaling factor is sufficient to make observed transcript counts comparable across cells, experiments, or platforms.

To illustrate this point, we first compared genes shared across MERFISH experiments with panels of different sizes. Total detected transcripts per cell decreased with increasing panel size (Fig. 1A), indicating that the same shared genes contribute fewer detected transcripts as panel size increases. We next compared log-transformed proportional expression of these shared genes between panels (Fig. 1B). If proportional expression were preserved across panels, genes would fall along the identity line (the line on which proportional expression is equal in both panels). Instead, many genes systematically deviated from this relationship, indicating that proportional expression is distorted between the larger and smaller gene panels. Similar distortions were also observed across independent imaging platforms (Plummer et al., 2025) (Fig. 1C; Additional File 2: Fig. S1 and Table S1) consistent with the principle of subcompositional incoherence, whereby relative abundances depend on the subset of features being analyzed rather than remaining invariant across subcompositions (Aitchison, 1986). This limitation is not unique to imaging-based ST. Across sequencing technologies, measured counts alone contain insufficient information to identify global transcript abundance, and normalization imposes assumptions about an unobserved quantity rather than estimating it directly (Nixon et al., 2025; McGovern & Silverman, 2026). When these assumptions are violated, DE analyses may incorrectly attribute changes in relative transcript composition to changes in absolute abundance, leading to biased effect-size estimates and mis-calibrated inference. These observations have motivated a class of scale-reliant inference methods that explicitly represent uncertainty in transcriptomic scale rather than fixing it through normalization. Rather than treating global scale as known following normalization, scale-aware approaches propagate uncertainty in absolute transcript abundances throughout downstream statistical inference.

**Figure 1.**
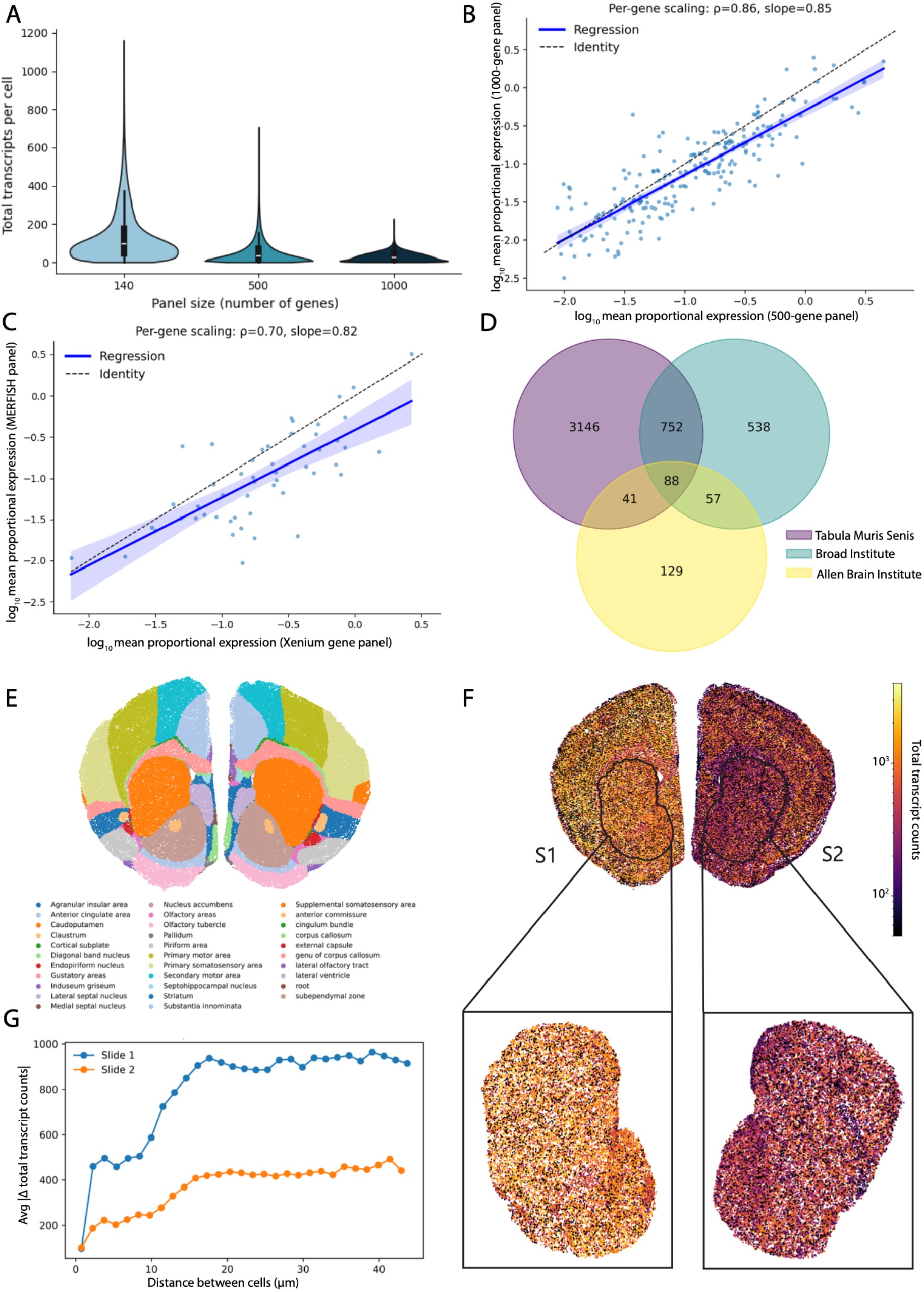
Spatial count-compositional structure in ST data. ST measurements exhibit both global scale variability and spatially structured compositional effects. Sequencing depth (scale) depends on panel design and assay efficiency. *(a–d) Global scale uncertainty*. (a)Total transcripts per cell computed using only genes shared across MERFISH panels (140, 500, 1000 genes) decrease with increasing panel size. (b) Comparison of log_10_ mean proportional expression for genes shared between the 500- and 1000-gene MERFISH panels. Deviations from the identity line indicate that proportional expression is not preserved across panel compositions. (140- vs 500- panel comparison shown in Supplementary Fig. S1.) (c). Comparison of log_10_ mean proportional expression for genes shared between MERFISH and Xenium platforms. Deviations from the identity line indicate that proportional expression is not preserved across platforms. (d) Overlap of differentially expressed genes of oligodendrocytes across three large single-cell aging atlases with limited concordance. *(e–g) Spatially structured compositional effects*. (e) Two matched coronal sections from biological replicates, colored by anatomical region, showing similar spatial organization. (f) Sections depicted in (e) colored by total transcript counts, revealing distinct global and local capture- depth patterns despite anatomical similarity. (g) Mean absolute differences in total transcript counts as a function of neighborhood radius. Differences increase with spatial distance, reflecting local compositional structure, while curves remain offset between sections, indicating persistent global scale differences.

Additionally, in some biological settings such as aging, total RNA abundance is not constant but can shift systematically across cell and tissue types, rendering technical and biological contributions to global scale inseparable. Several studies have reported transcriptome-wide reductions in transcriptional output with age (Stoeger et al., 2022; Uemura, 1980), including evidence that this decline may arise in part from stalling of RNA Polymerase II (Gyenis et al., 2023). Consistent with these challenges in identifiability, comparison of DE results across three large single-cell mouse brain aging atlases published by the Tabula Muris Senis consortium (2020), Broad Institute (Ximerakis et al., 2019), and the Allen Institute (Jin et al., 2024) revealed limited overlap in gene-level signatures despite substantial numbers of significant genes in each dataset (Fig. 1D; example shown for oligodendrocytes). Although many factors contribute to cross-study variability, these observations illustrate the broader reproducibility issues that arise when changes in absolute transcript abundance cannot be distinguished from changes in relative transcript composition.

Together, these observations indicate that global scale variability is not merely a technical nuisance, but a fundamental source of uncertainty in imaging-based ST. This uncertainty extends beyond DE to downstream analyses such as imputation and reference-based cell type annotation (B. Li et al., 2025; Shen et al., 2025; Zhou et al., 2025), which often implicitly assume that proportional expression of marker genes is preserved across experiments following normalization. The observed distortions in proportional expression across panels and platforms (Fig. 1B-C; Additional File 2: Fig S1) suggests that this assumption may not always hold, potentially contributing to unstable cross-platform mapping and reference alignment. Such observations call for statistical frameworks based on the principles of scale reliant inference, in which uncertainty in global transcript abundance is modeled explicitly.

#### Compositional uncertainty

The second component of count-compositional uncertainty arises from local transcript composition. Within each cell, transcript abundance is represented as a composition—a vector of positive transcript proportions that sum to one. Consequently, changes in the relative abundance of one transcript necessarily alter the relative abundance of the remaining transcripts. Even between closely matched coronal sections from biological replicates, transcript depth exhibits strong slide-specific spatial structure. Although sections share similar anatomical organization, total transcript counts differ both globally and locally, with smooth gradients evident within matched regions, such as the caudoputamen and the nucleus accumbens (Fig. 1E–F). Notably, these gradients differ in direction and magnitude between slides despite similar anatomy, suggesting that transcript depth is influenced by slide-specific technical variation in addition to biological structure.

This structure can be quantified by examining transcript depth across spatial neighborhoods. At small spatial scales, neighboring cells show relatively stable transcript depth, while differences increase with spatial distance, reflecting local compositional structure across tissue space (Fig. 1G). Importantly, sections remain offset across all spatial scales, indicating persistent global scale differences superimposed on local variation.

These patterns might suggest that modeling spatial correlation is sufficient to account for structured variability. However, not all structured dependence in imaging-based ST is spatial. Technical factors such as imaging rounds and fluorescence channels can introduce gene-specific covariance unrelated to tissue proximity (Martin et al., 2025), while biologically meaningful clustering may itself be spatially localized (Sarwar et al., 2025). As a result, explicitly attributing local structure to spatial correlation may partially absorb true biological signals.

Together, these observations indicate that imaging-based ST measurements contain two complementary sources of uncertainty: uncertainty in total transcript abundance and uncertainty arising from local transcript composition. The framework presented below represents these components separately, allowing uncertainty in both global scale and relative composition to be propagated throughout differential-expression inference.

### ALDEx model overview

To address the compositional and scale uncertainties described above, we adapt the Bayesian partially identified ALDEx3 mixed-effects framework (McGovern and Silverman, 2026) to imaging-based ST data. In this formulation, observed transcript counts are treated as count-compositional measurements: counts reflect both the relative composition of transcripts within each cell and uncertainty in total transcript abundance across cells and experimental conditions. The framework represents these two sources of uncertainty separately and propagates both through DE inference.

#### Model decomposition

Let *W* denote the latent transcript abundance matrix, with rows corresponding to genes and columns corresponding to cells. We index cells by *n*, such that *W*·_*n*_ denotes the vector of latent transcript abundances across all genes in cell, where the dot indicates all genes. We decompose log-abundance for each cell into compositional and global scale components,

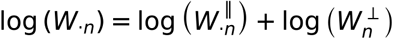

where 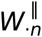 represents relative transcript composition across genes within cell *n*, and 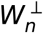 is a latent global scale factor shared across all genes in that cell. This decomposition separates variation arising from relative transcript composition from uncertainty in total RNA abundance.

#### Measurement model

We represent the compositional component using the ALDEx Monte Carlo framework (McGovern and Silverman, 2026). Conditional on latent transcript composition *p*_*n*_, observed counts are modeled as

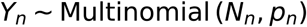

with a Dirichlet prior placed on *p*_*n*_, yielding a posterior distribution over compositions (Methods). Monte Carlo samples from this posterior define realizations of the compositional component 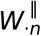, capturing uncertainty arising from finite sampling depth under the constraint that counts sum to the total number of detected transcripts in each cell.

#### Structured scale framework

Unlike transcript composition, global transcript abundance is not identifiable from the observed counts alone. Thus, within the Bayesian partially identified framework, uncertainty in global scale is represented through a prior distribution rather than estimated directly from the data. Specifically, for each cell *n*, the latent scale component is assigned the prior

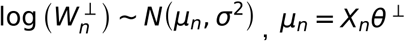

where *X*_*n*_ denotes the row of the design matrix corresponding to cell *n,θ*^⊥^ represents predictor-associated effects on global scale and *σ*^2^ specifies residual uncertainty in transcript abundance not explained by the predictors.

This prior is not intended to recover a single estimate of global transcript abundance. Instead, it propagates plausible uncertainty about scale throughout downstream inference, allowing transcriptome-wide shifts in RNA abundance to be represented separately from gene-specific compositional changes. When applicable, prior information may be obtained from previous experimental studies or biological knowledge. For example, evidence for transcriptome-wide reductions in RNA abundance during aging (Stoeger et al., 2022; Gyenis et al., 2023) motivates the asymmetric scale prior used in our aging analyses. Such priors constrain the direction of the scale effects while retaining uncertainty in their magnitude (Nixon et al., 2025). Practical implementation details, including mixed-effects inference engines and computational scaling considerations, are described in Methods.

#### Key implications

In this framework, gene-level effect estimates arise from the compositional model, while uncertainty about global transcript abundance is propagated through the Bayesian scale priort. Because the scale term is shared across genes, it does not directly alter gene rankings but instead modulates statistical confidence in gene-specific effects. This allows transcriptome-wide shifts in RNA abundance to be accounted for without conflating them with gene-specific DE.

### Benchmarking performance

To evaluate whether scale-aware DE models remain calibrated under realistic spatial transcriptomic measurement constraints, we constructed a simulation framework based on a MERFISH aging dataset acquired by our group (Fig 2A; Methods), comprising paired young and old brain hemi sections per run across multiple biological replicates. Simulations preserve the original count structure, batch organization, and compositional coupling while enabling controlled manipulation of signal and sample size. We benchmarked ALDEx against widely used DE approaches commonly applied to replicated ST datasets.

**Figure 2.**
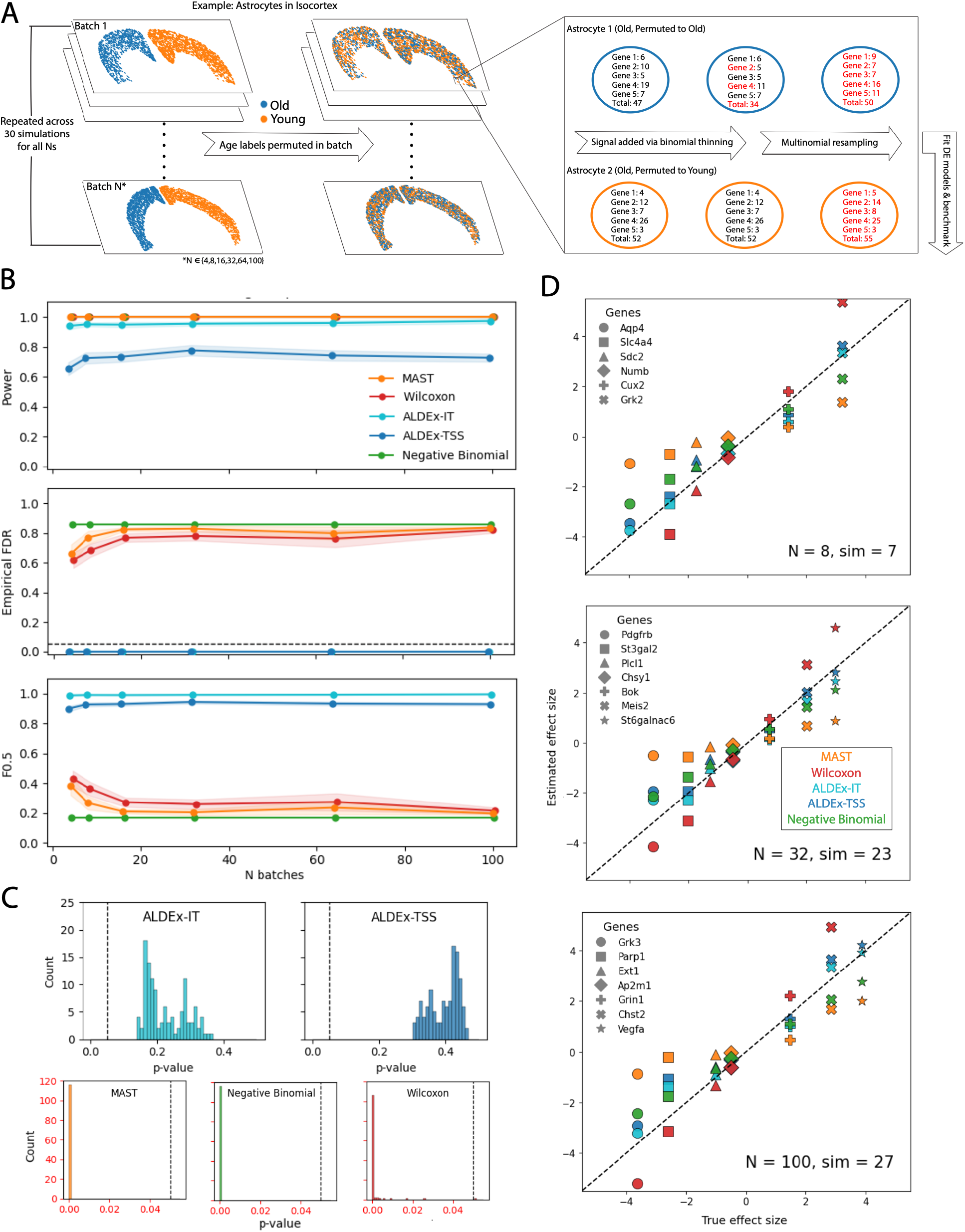
Benchmarking of DE methods on simulated data. (a) Overview of the simulation strategy. Observed ST data with original age labels (young/old) were permuted within their corresponding batch to generate exchangeable null datasets. Example cells illustrate reassignment of labels following permutation. (b) Benchmarking performance across increasing sample size (number of batches), showing empirical FDR, power, PPV, and F0.5 score. Points and lines represent mean performance across simulations, with shaded regions indicating 95% confidence intervals. (c) Distribution of null p-values at a representative sample size (N = 32). Histograms are shown for each method using genes without injected signals. The dashed line indicates the nominal significance threshold (p = 0.05). (d) Recovery of true effect sizes across methods. Estimated coefficients are plotted against true injected effects for a representative subset of genes spanning the range of effect sizes. Panels correspond to increasing sample sizes (N = 8, 32, 100) and show randomly selected example simulations. The diagonal line indicates perfect recovery.

Null datasets were generated by permuting age labels within batches of the target cell type in a target region (e.g. astrocytes in the isocortex), preserving spatial data structure while removing association with expression (Fig 2a; Methods). Controlled gene-specific signal was introduced using binomial thinning, with differential effects assigned asymmetrically (80% downregulated and 20% upregulated in the old group) to reflect expected global transcriptional decline with aging. Thinning imposes the desired log2 fold-change pattern while preserving the underlying count structure (Gerard, 2020), followed by multinomial resampling to restore finite-count compositional measurement constraints. Performance was evaluated across increasing study sizes by resampling batch units to generate datasets ranging from 4 to 100 batches.

We evaluated two ALDEx formulations: an agnostic structured scale model (ALDEx-TSS) and an informed structured scale model in which age is allowed to induce a global shift in transcript abundance through a non-zero prior on scale effects (ALDEx-IT). For comparison, we included widely used cell-level DE methods, including MAST (hurdle model framework) (Finak et al., 2015), Wilcoxon rank-based testing as implemented in Scanpy (a common default in single-cell and ST workflows) (Wolf et al., 2018), and negative binomial generalized linear models (Hafemeister and Satija, 2019). These methods represent standard analytical approaches currently used in the field (Kalantari-Dehaghi et al., 2025; Wang et al., 2026) and provide a baseline for evaluating how explicitly modeling compositional constraints and global transcript abundance affects inference.

Across all sample-size scenarios, both ALDEx formulations remained well calibrated (Fig. 2B; Additional File 3), with empirical false discovery rates (FDR) effectively at zero and precision near unity. The agnostic TSS model was more conservative, yielding moderate but stable power, whereas the informed structured scale model substantially improved sensitivity while preserving strong error control (Fig. 2B). These results indicate that incorporating biologically grounded scale priors improves recovery of true signal without compromising calibration. This conservative calibration is further reflected in the distribution of p-values under null simulations, which are skewed toward larger values (Fig. 2C), reflecting conservative inference under compositional uncertainty.

In contrast, methods that do not explicitly model compositional constraints exhibited systematic miscalibration. The Wilcoxon test showed severe inflation of false discoveries, with empirical FDR increasing substantially with sample size (Fig. 2B; Additional File 3). Although power remained maximal, positive predictive value and F0.5 (a metric emphasizing precision over recall) declined in parallel, indicating that increased sample size amplified false positives rather than improving inference. MAST and negative binomial models displayed more moderate but persistent deviations from nominal behavior: both achieved near-maximal power across all sample sizes, but empirical FDR remained above the nominal threshold and did not improve with increasing batch number (Fig. 2B). Correspondingly, their null p-value distributions show excess mass at small p-values relative to uniform expectations (Fig. 2C), consistent with anti-conservative behavior and inflated false-positive rates.

Examination of effect-size recovery across representative simulations further illustrates gene-level behavior across methods (Fig. 2D; Additional File 3). Estimated coefficients from ALDEx, negative binomial, and Wilcoxon approaches all remained strongly correlated with the true simulated effect sizes across sample sizes, indicating that each method was capable of recovering the underlying injected signal. In contrast, MAST exhibited substantially greater dispersion and attenuation of effect estimates. While effect-size recovery was broadly similar among the best-performing methods, their inferential properties differed markedly: only the ALDEx formulations consistently maintained strong control of FDR across simulation scenarios (Fig. 2B–C).

Together, these results demonstrate that explicit modeling of compositional constraints and global scale substantially improves the reliability of DE inference. By propagating uncertainty in both local composition and global transcript abundance, ALDEx maintains strong calibration and false-discovery control across simulation scenarios while preserving accurate recovery of underlying biological effects.

### Cross-study reproducibility

To demonstrate the utility of our scale-aware DE framework, we first analyzed a MERFISH aging dataset generated by our group, comprising young (2-month-old) and aged (22-month-old) mouse brains profiled using a custom 500-gene panel (Fig. 3A). Following preprocessing, cell-type annotation, and anatomical registration (Methods), DE was analyzed across 34 non-neuronal cell type × region strata using the final informed scale-aware ALDEx model, which propagates uncertainty in both transcript composition and global RNA abundance. Across these strata, 446 significant aging-associated DE results were identified, corresponding to 130 unique genes, with most significant effects reflecting reduced expression with age (Fig. 3B–C; Additional File 4).

**Figure 3.**
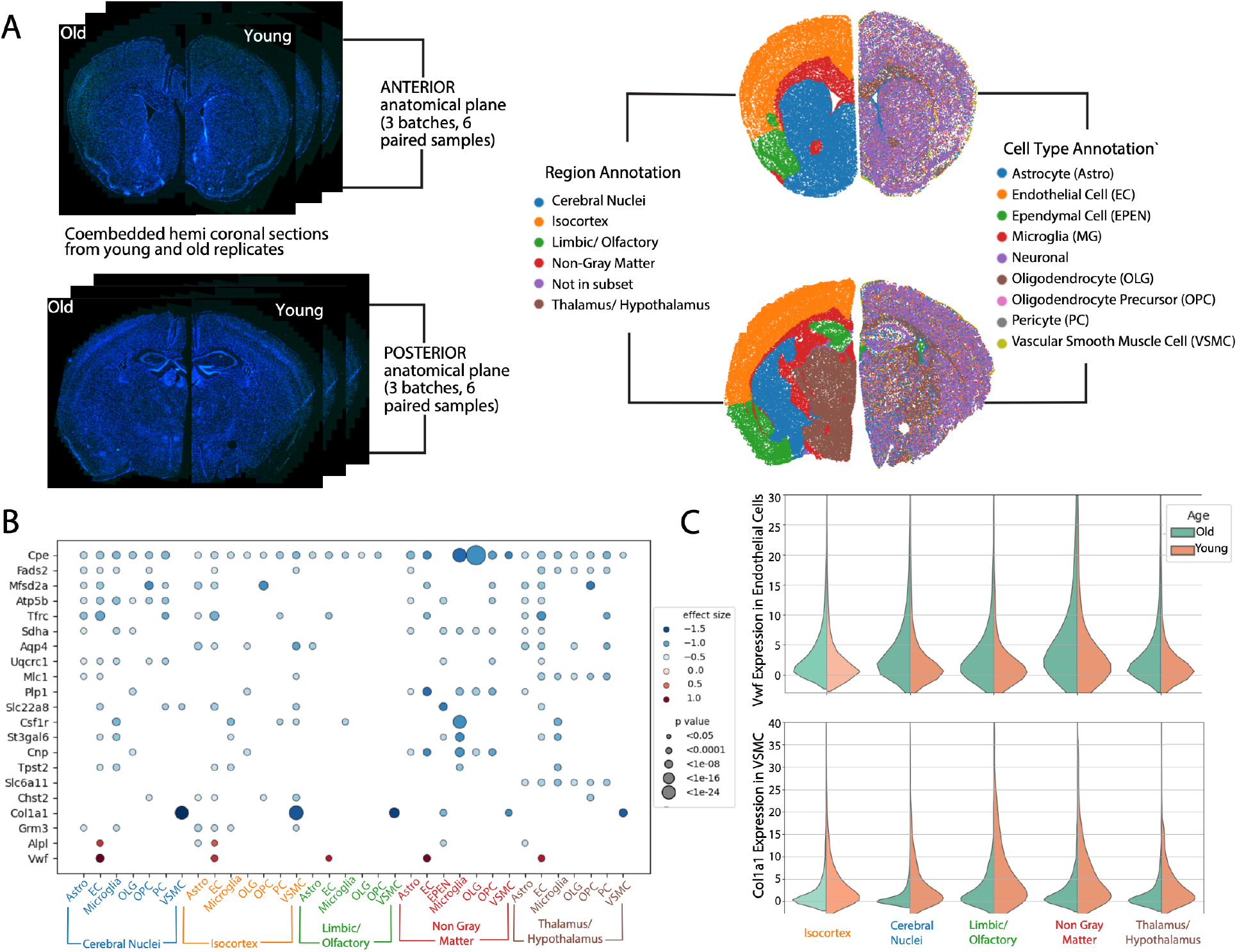
Aging-associated transcriptional programs across spatially resolved glial and vascular populations. (a) Overview of experimental design and spatial annotation. DAPI images show co-embedded young and old hemi-coronal brain sections processed across six batches, spanning anterior and posterior anatomical planes. Representative paired sections from each plane are shown. Spatial maps illustrate cells colored by broad anatomical regions (left) and by cell type annotations (right). (b) DE across cell type × region strata. Dot plot of representative genes, with color indicating effect size (log fold change; red = higher in old, blue = lower in old) and dot size proportional to statistical significance (−log_10_ adjusted p-value). (c) Representative expression distributions supporting DE results. Split violin plots show raw expression of *Vwf* in endothelial cells and *Col1a1* in vascular smooth muscle cells across regions, comparing young and old samples. Observed shifts in expression are consistent with inferred differential effects.

To determine whether these signals reflected reproducible biological processes rather than study-specific effects, we next evaluated their consistency across independent spatial transcriptomic datasets spanning different experimental designs and levels of anatomical correspondence (Methods).

#### Validation in a platform-matched dataset

We first evaluated reproducibility using the coronal subset of a mouse brain aging MERFISH dataset (Sun et al., 2025), which profiles gene expression across a continuous age range (3–34 months) in anatomically matched sections. In the original study, aging-associated changes were characterized using complementary analyses including pseudobulk Spearman correlation, trajectory-based classification of lifespan expression dynamics, and spatial aging clocks trained on spatially smoothed transcriptomes. These analyses provided an opportunity to evaluate whether scale-aware DE signals converge with independently derived aging-associated biological programs.

To enable direct comparison between methods, we harmonized the Sun dataset to a binary young-versus-old design and analyzed both the Sun and our primary datasets using ALDEx and an adapted version of the published Sun pseudobulk framework (Methods; SAdditional Files 4-5). Reproducibility was then assessed by comparing gene-level aging signals between datasets within matched cell type × region strata, separately for each method (ALDEx-to-ALDEx and pseudobulk-to-pseudobulk comparisons).

We first evaluated the reproducibility of aging-associated effect directions (Fig. 4A). Across shared cortical non-neuronal cell types in the isocortex, ALDEx preserved the direction of aging-associated effects for the majority of intersecting genes in the two datasets, with discordant calls limited to a small subset. In contrast, the adapted pseudobulk baseline produced substantially more reversals of effect direction across endothelial, immune, oligodendrocyte-lineage, and vascular-associated populations. These results indicate that scale-aware modeling improves the stability of effect direction across studies.

**Figure 4.**
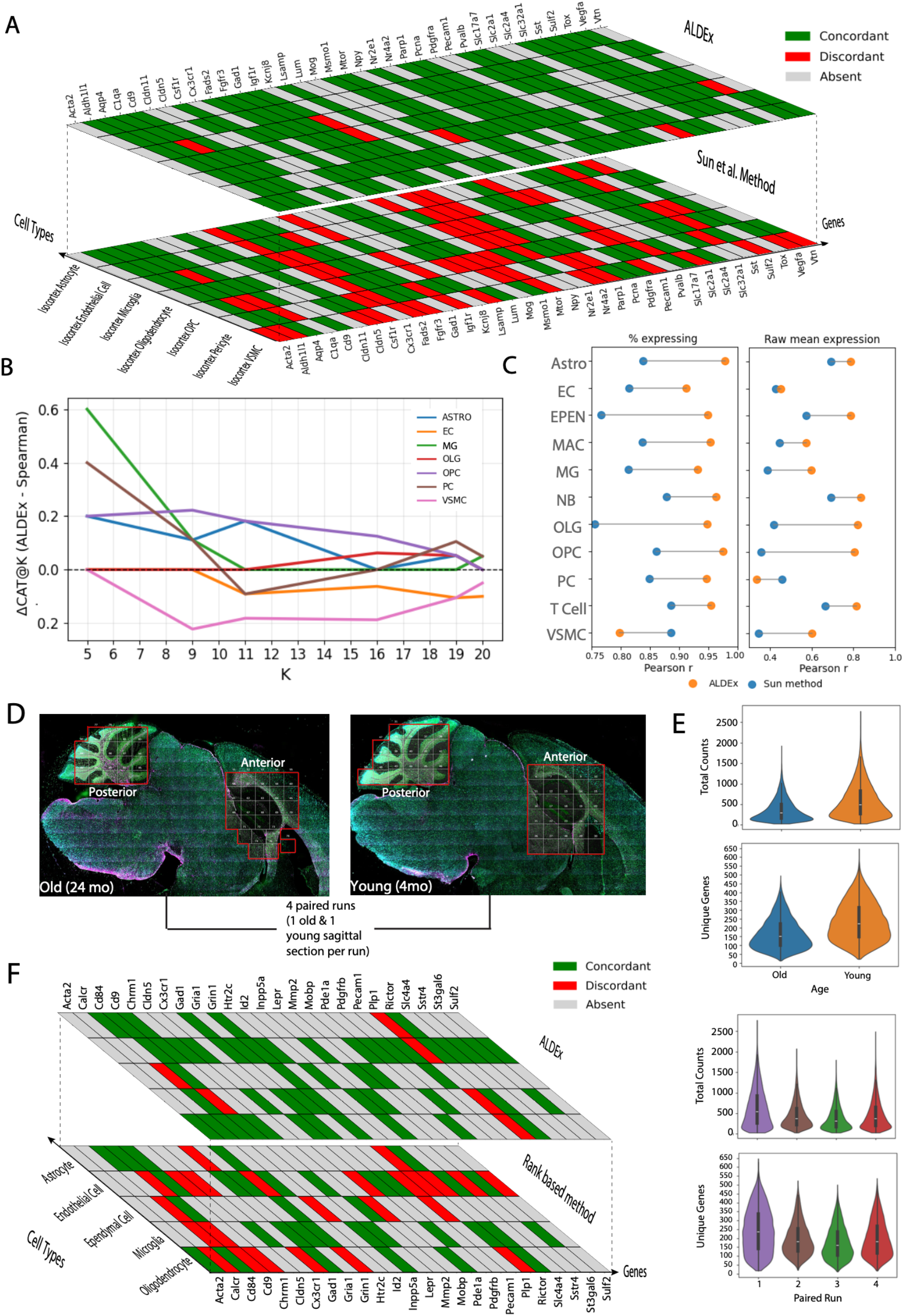
Cross-study reproducibility of scale-aware DE signals. *(a–c) Within platform comparison*. (a) Heatmaps showing sign concordance of aging-associated DE effects between our MERFISH aging dataset and the Sun et al. coronal aging dataset across matched non-neuronal populations for ALDEx and the adapted Spearman/pseudobulk framework. (b) ΔCAT@K analysis comparing overlap among top-ranked aging-associated genes between datasets across a range of K values. Positive values indicate improved overlap under the ALDEx framework relative to the adapted Spearman/pseudobulk baseline. (c) Comparison of inferred aging-associated effect sizes with corresponding changes in unnormalized transcript abundance metrics using matched extreme-age cohorts from the Sun dataset (3.4–5.4 months versus 30.9–34.5 months). Agreement was evaluated using Pearson correlation between DE effect estimates from continuous-age ALDEx or the published continuous-age Spearman framework and observed shifts in mean raw transcript abundance or expression frequency (fraction of expressing cells) between young and old groups across matched non-neuronal populations. *(d-f) Between platform comparison*. (d) Overview and experimental design of SenNet CosMx dataset, consisting of 4 paired runs/ batches of sagittal sections from one young (4mo) and one old (24 mo) female mouse, each containing one anterior (hippocampus) and one posterior (cerebellum) region of interest. (e) Split violin plots showing average total counts (upper panel) and average number of unique genes expressed per cell (lower panel) stratified by age (left) and paired run (right). (f) Heatmaps showing sign concordance of aging-associated DE effects between the primary MERFISH aging dataset and the independent CosMx SenNet aging dataset across matched non-neuronal populations for ALDEx and the adapted rank-based pseudobulk framework. Concordance was computed on intersecting genes within matched cell types.

We next asked whether the genes prioritized most strongly by each method were reproducible across datasets. Across cortical glial and vascular populations, ALDEx showed greater overlap among the highest-ranked aging-associated genes than the adapted pseudobulk Spearman framework (Fig. 4B). This advantage was quantified using differences in concordance at the top (ΔCAT@K), a metric that compares overlap among the top K ranked genes between methods; positive values indicate greater overlap under ALDEx. The largest improvements were observed in cortical astrocytes and immune populations and were most pronounced at smaller values of K, indicating that ALDEx more consistently prioritized the strongest and most reproducible aging-associated signals. Differences between methods diminished at larger K values as ranked gene lists became increasingly similar. Effect-size concordance showed a similar but weaker pattern (Additional File 2: Table S2). ALDEx achieved comparable or improved Pearson and Spearman correlations in most matched comparisons.

To further evaluate the biological consistency of the inferred aging signals, we performed supplementary analyses using the full Sun coronal dataset with age modeled as a continuous predictor (Additional File 5). Continuous-age ALDEx showed modestly higher agreement with the published trajectory analysis and greater overlap precision with aging-clock-associated Gene Ontology Biological Process (GO-BP) enrichments despite identifying fewer significant genes overall (Additional File 2: Fig. S2A-B). Notably, pathways associated with transcriptional regulation, RNA metabolism, and RNA polymerase II activity were consistently classified as decreasing with age across trajectory, aging-clock, and ALDEx analyses, whereas the published Spearman framework identified a slight majority of genes as increasing (Sun et al., 2025; Additional File 5). Comparison against raw transcript abundance and expression frequency further showed that ALDEx effect estimates more closely reflected observed age-associated changes, particularly for gene detection frequencies (Fig. 4C; Additional File 2: Fig. S2C). Together, these results suggest that scale-aware modeling more effectively captures coordinated age-associated transcriptional decline within the Sun dataset.

#### Validation in a cross-platform dataset

To assess the reproducibility of ALDEx-derived aging-associated signals across independent imaging-based ST platforms, we reanalyzed a publicly available CosMx Spatial Molecular Imaging (SMI) mouse brain aging dataset generated through the SenNet Consortium and recently described by Carver et al. (2026) . The dataset consisted of paired experimental runs comprising sagittal brain sections from four young (4 month) and four aged (24 month) female mice, profiled using the 950-plex CosMx Mouse Neuroscience panel supplemented with a custom 50-gene-aging- and senescence-associated probe panel. Tiled anterior and posterior sagittal regions of interest corresponded primarily to hippocampal/corticolimbic and cerebellar areas, respectively (Fig 4D-E). The CosMx dataset provided an independent cross-platform comparison using a distinct imaging chemistry and sagittal sampling strategy while retaining a similarly paired binary aging design and greater overlap with the primary gene panel.

We performed independent preprocessing and cell-type assignment using the raw data available through the SenNet Data Portal at the time these analyses were initiated. (Methods; Additional File 2: Fig S3A-C). Downstream DE analyses were harmonized with our dataset as closely as possible using matched ALDEx modeling parameters and mixed-effects model structure. Both ALDEx and the adapted pseudobulk framework were then applied to the CosMx data using the same cross-study comparison strategy described above, albeit at the cell type level collapsed across anatomical regions (Methods; Additional File 6).

ALDEx generally showed improved directional agreement, ranking concordance, and effect-size reproducibility relative to the adapted rank-based pseudobulk framework, despite substantial differences in platform chemistry, section orientation, and sex between studies (Fig 4F; Additional File 2 Fig. S3D and Table S3; Additional File 6).

### Recurrent aging-associated programs across non-neuronal populations

Having established the reproducibility of the inferred aging-associated signals across independent datasets, we next examined the biological programs underlying these recurrent changes. We focused on non-neuronal populations, including glial and vascular cell types, which play central roles in brain homeostasis yet remain comparatively understudied in aging research. To facilitate biological interpretation, Gene Ontology (GO) and Kyoto Encyclopedia of Genes and Genomes (KEGG) pathway annotations associated with significant genes from our MERFISH dataset were aggregated into broader functional modules that summarize recurrent biological themes rather than serving as formal pathway-enrichment statistics (Methods; Additional File 6). This analysis identified a small number of recurring biological themes spanning neurovascular support, membrane transport and homeostasis, extracellular matrix remodeling, and growth and survival signaling (Fig. 5A-C).

**Figure 5.**
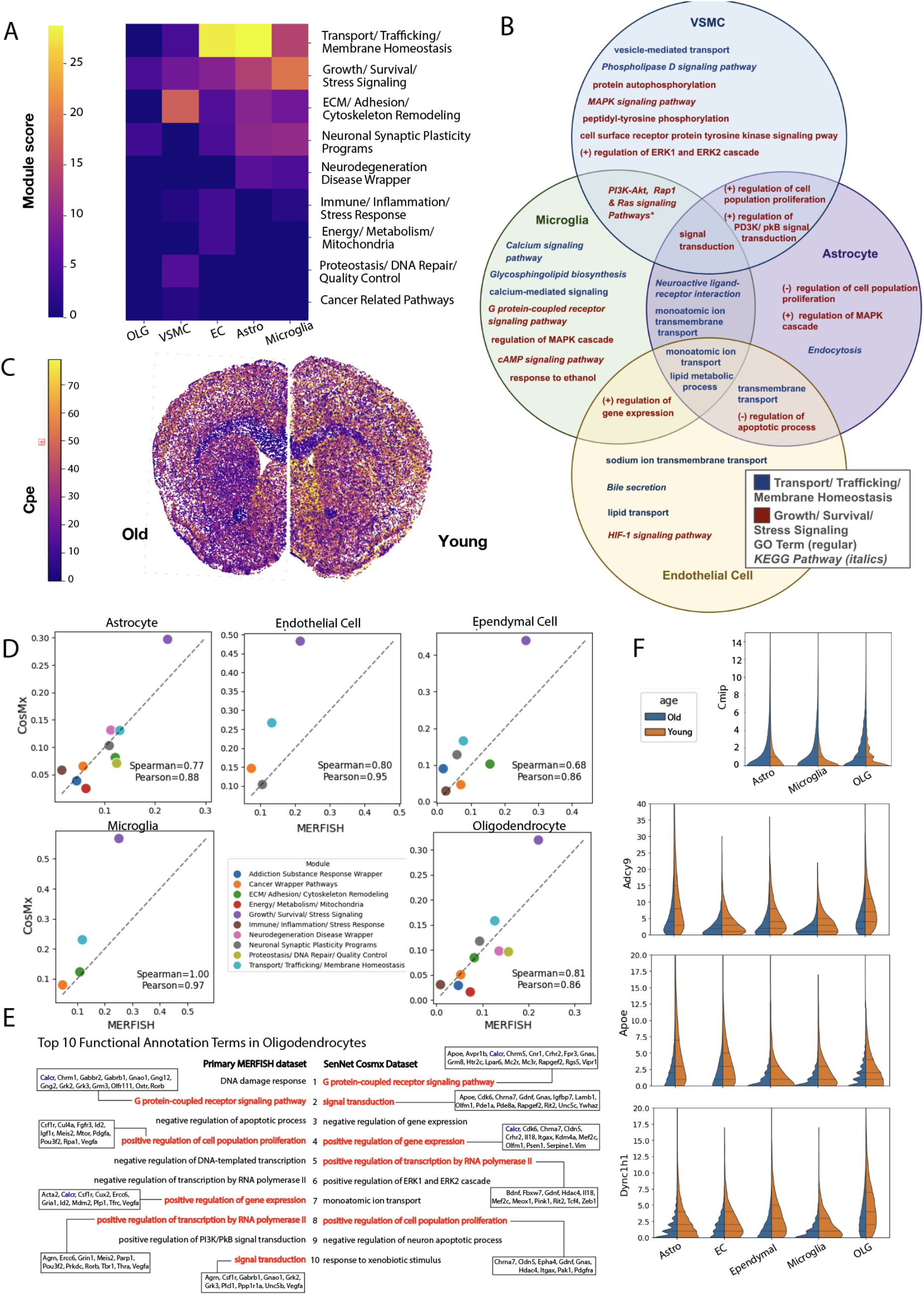
Recurrent signatures of senescence in non-neuronal populations across datasets. (a) Functional module summary of aging-associated programs. Heatmap of module scores across cell types, where values represent the relative contribution of aggregated functional annotations within each module. (b) Overlap of functional annotations across cell types. Venn diagram showing shared and cell type–specific terms across astrocytes, endothelial cells, microglia, and vascular smooth muscle cells within the top modules, highlighting common biological processes affected by aging. (c) Example paired young and old sections colored by raw counts of *Cpe* across all cells, illustrating global differences in expression consistent with model estimates. (d) Comparison of module-level functional signatures between the primary MERFISH dataset and the CosMx dataset. Each point represents a functional module, with module scores computed from weighted GO and KEGG annotation contributions aggregated within matched cell types. Dashed lines indicate equality between datasets. Spearman and Pearson correlations summarize concordance of module-level aging signatures across platforms. (e) Annotation-level comparison of aging-associated functional programs in oligodendrocytes. Top-ranked GO and KEGG annotations from each dataset are shown alongside their contributing genes, with connecting lines indicating annotations recovered in both studies. (f) Representative examples of aging-associated genes identified in the independent CosMx SenNet aging dataset. Violin plots show raw transcript abundance distributions in young and aged cells for selected genes exhibiting consistent decreases or increases across multiple glial populations. Additional examples are provided in Supplementary Fig. 3.

We next asked whether these same higher-order programs could be recovered in the independent CosMx dataset despite substantial differences in platform chemistry, anatomical sampling, panel composition, and sex. To enable direct comparison across platforms, module-level analyses were repeated using only functional annotations shared between datasets (Methods; Additional File 6). Although overlap among individual significant genes was limited, the independently derived functional module summaries were concordant across matched non-neuronal populations (Fig. 5D-E). Agreement was particularly notable given the distinct characteristics of the two studies, suggesting that higher order biological themes are substantially more reproducible across datasets than individual aging-associated genes.

One of the most consistent signals involved genes associated with transport, trafficking, and membrane homeostasis, particularly lipid transport and transmembrane exchange. In our dataset, this module was prominent across astrocytes and endothelial cells (Fig. 5A) and was supported by recurrent genes including Mfsd2a, Fads2, Slc2a1, and Tfrc. These genes participate in the transport and metabolism of essential lipids and metabolites at the blood–brain barrier and within glial metabolic networks (Panov et al., 2014). For example, Mfsd2a mediates transport of docosahexaenoic acid (DHA) across the blood–brain barrier (Nguyen et al., 2014; Z. Wang et al., 2020), while Fads2 catalyzes a rate-limiting step in the synthesis of long-chain polyunsaturated fatty acids (Nakamura et al., 2004). Astrocytes are known to regulate lipid metabolism and membrane composition within the neurovascular unit (Bolaños and Almeida, 2026; Moore et al., 1991), and endothelial cells control molecular transport across the blood– brain barrier (Xiang et al., 2025). The convergence of lipid metabolic and transport-related signals across these cell types identified transport and membrane homeostasis as a prominent aging-associated program. Consistent with this observation, many of the associated functional annotations were shared across astrocytes, endothelial cells, vascular smooth muscle cells, and microglia (Fig. 5B).

Similar patterns were observed in the independent CosMx dataset, where transport and membrane homeostasis emerged as one of the highest-ranked aging-associated modules. Although fewer functional annotations contributed to this module in the cross-platform comparison framework, its relative ranking remained highly consistent across studies (Fig. 5D). Representative CosMx genes included Dync1h1, which encodes the heavy chain of cytoplasmic dynein and plays a central role in intracellular transport (Braunstein et al., 2010), and Apoe, a key regulator of lipid metabolism and one of the strongest genetic modifiers of Alzheimer’s disease risk (Shinohara et al., 2016) (Fig. 5F). Together, these observations suggest that age-associated alterations in intracellular trafficking, lipid handling, and membrane homeostasis constitute a recurrent biological program across glial and vascular populations in the aging brain.

A second recurring pattern involved genes associated with growth, survival, and stress signaling, including pathways related to MAPK and PI3K–AKT signaling. Notably, the neuropeptide-processing enzyme Cpe (carboxypeptidase E) was detected across 32 of 34 cell type–region strata (Fig. 3B; Fig 5C). CPE has been implicated in ERK/AKT signaling, neuronal survival, and BDNF maturation (Cheng et al., 2013; Fan et al., 2023; Liu et al., 2023). The widespread appearance of Cpe alongside recurrent decreases in MAPK- and PI3K-related pathways identified growth and survival signaling as a second major aging-associated program.

The prominence of this signaling program was even more apparent in the independent CosMx dataset, where growth, survival, and stress signaling represented the highest-ranked functional module across all major non-neuronal populations (Fig. 5D). Representative CosMx genes included Ywhaz, a component of insulin and IGF-associated signaling pathways implicated in longevity, cellular survival, and neurodegeneration (Wang, 2012), and Adcy9, which encodes an adenylyl cyclase responsible for cyclic AMP production and has previously been reported to decline in the aged hippocampus, where reduced cAMP signaling has been linked to age-associated impairments in learning and memory (Mons et al., 2004) (Fig 5F).The concordance of both module-level signatures and recurrent signaling-associated genes across datasets points to coordinated age-associated reductions in trophic, metabolic, and stress-response pathways across multiple non-neuronal populations.

Extracellular matrix (ECM) organization, cell adhesion, and cytoskeletal remodeling represented a third major theme, particularly within vascular-associated populations. These signals were strongest in vascular smooth muscle cells (Fig 5A), and were supported by genes including Col1a1, Pdgfrb, Tgfbr3, and Lama1, which are involved in extracellular matrix production, vascular structure, and cell migration (Fig 3C) (S. Li et al., 2000; Makihara et al., 2015; Massagué and Sheppard, 2023).

Although individual ECM-associated genes were not strongly shared across datasets, extracellular matrix remodeling remained concordant at the module level in the CosMx comparison (Fig. 5D). Interpretation of gene-level overlap was further limited by the absence of a sufficiently large vascular smooth muscle cell population in the CosMx dataset, likely reflecting differences in anatomical sampling. Nevertheless, the persistence of extracellular-matrix-associated signatures at the module level suggests that age-associated changes in vascular structure and extracellular-matrix organization are more consistently recovered at the level of biological processes than at the level of individual genes.

Taken together, these analyses identified recurrent aging-associated signatures involving transport and membrane homeostasis, growth and survival signaling, and extracellular matrix remodeling across multiple non-neuronal populations. Although the specific genes contributing to these signatures differed substantially between datasets, concordance at the level of functional modules remained strong. These findings indicate that higher-order biological programs are more consistently identified across independent spatial transcriptomic studies than individual gene-level signals.

## Discussion

Spatial transcriptomics (ST) has rapidly expanded the scope of questions that can be addressed in intact tissues, yet DE analysis remains a methodological challenge. Existing methodological advances in ST address complementary statistical challenges. Spatial correlation-based DE approaches are primarily designed for within-sample inference, where expression is modeled against spatially varying covariates such as anatomical region, local cellular neighborhood, or distance-dependent effects (Cable et al., 2022; Mason et al., 2024; Ospina et al., 2024; Vasconcelos et al., 2026; Wang et al., 2026). In contrast, many biological studies, including aging, development, and treatment-response experiments, are fundamentally replicated between-sample comparisons (Cao et al., 2024; Kukreja et al., 2026; S. Li et al., 2025). In these settings, investigators typically rely on cell-level or pseudobulk workflows adapted from single-cell RNA-seq despite the distinctive compositional and spatial characteristics of ST measurements (Cui et al, 2025; Kiviaho et al., 2024; Sun et al., 2025). Here, we addressed this gap by extending the ALDEx framework to jointly account for compositional uncertainty and variation in global transcript abundance while retaining cell-level resolution and support for replicated experimental designs, which help to identify converging biological pathways upon aging across different ST datasets.

Simulated benchmarking experiments demonstrated that explicitly modeling these sources of uncertainty improved calibration under realistic ST measurement constraints. Across increasing sample sizes, both ALDEx formulations maintained strong control of false discoveries, whereas methods that did not explicitly account for compositional structure exhibited consistent or increasing miscalibration. Notably, the inflation of FDR was not remediated by larger sample sizes, highlighting that additional data do not necessarily improve inference when underlying measurement assumptions are mis-specified. Rather, increasing sample size may amplify confidence in biased estimates. The informed structured-scale model further improved sensitivity while maintaining strong error control, suggesting that biologically motivated assumptions about transcriptome-wide changes can increase power without sacrificing calibration.

The benchmarking framework was similarly designed around the replicated between-sample setting considered here. Accordingly, we did not benchmark methods intended for within-sample analyses, as doing so would have required substantially changing the underlying biological question. Instead, we focused on compositional and scale uncertainty while using independent real-world datasets to evaluate reproducibility and biological coherence under realistic experimental conditions. The external validation analyses supported both the ALDEx framework and the biological conclusions it produced. In the Sun et al. (2025) aging dataset, ALDEx consistently improved sign and rank concordance relative to the adapted pseudobulk Spearman framework, indicating more reproducible recovery of aging-associated signals across studies. ALDEx effect sizes also showed stronger agreement with independent measures of transcriptional change, including raw transcript abundance and expression frequency. The latter is particularly notable because expression frequency is a cell-level feature that is largely lost during pseudobulk aggregation. While pseudobulk restores replicate-level inference (Squair et al., 2021; Tung et al., 2017), it does so by collapsing cellular heterogeneity. In ST, where cell-level and spatial variation are often themselves of interest, preserving these features may improve recovery of biologically meaningful aging-associated signals.

The independent CosMx dataset provided an especially stringent test of biological reproducibility because it extended validation beyond independent cohorts to an independent ST technology. The primary study used CosMx together with complementary spatial and experimental approaches to identify senescence- and disease-associated microglial states concentrated in aged white matter (Carver et al., 2026). Our independent reanalysis addressed a distinct methodological question by evaluating the reproducibility of aging-associated differential-expression signals across platforms and across multiple non-neuronal populations. While overlap among significant genes was limited, concordance remained strong at the level of biological programs. Despite substantial differences in platform chemistry, anatomical sampling, panel composition, and sex, both the primary MERFISH and independent CosMx datasets repeatedly implicated processes related to cellular signaling, intracellular transport, membrane homeostasis, extracellular matrix organization, and maintenance of neurovascular support functions. Together, these findings suggest that higher-order aging-associated pathways may be more reproducible across studies than individual gene-level signatures and highlight the value of pathway-level interpretation when comparing independent spatial transcriptomic datasets.

Many of the recurrent programs identified by ALDEx converge on functions required for maintenance of the neurovascular unit. Transport and membrane-homeostasis signatures were recovered across astrocytes, endothelial cells, vascular smooth muscle cells, and microglia, suggesting that age-associated changes in lipid handling and membrane transport extend across multiple components of the neurovascular unit (M. Li et al., 2026; Cunnane et al., 2012; Zehr et al., 2018). Likewise, the widespread appearance of Cpe together with recurrent decreases in MAPK- and PI3K–AKT-associated pathways points to reduced trophic and stress-response signaling across glial and vascular populations (Cheng et al., 2013; Fan et al., 2023; Liu et al., 2023). Reproducible extracellular-matrix signatures provide a complementary indication of age-associated vascular remodeling (Elmarasi et al., 2024; Gao et al., 2020). Collectively, these observations suggest coordinated remodeling of interconnected systems responsible for neurovascular integrity, metabolic support, and trophic maintenance in the aging brain, and are consistent with broader models of declining cellular responsiveness and adaptive capacity during aging (López-Otín et al., 2023; Melo Dos Santos et al., 2024).

While the reproducibility observed across independent datasets supports the validity of the proposed framework, there are potential avenues for future refinement. Our primary objective was to address replicated between-sample differential-expression studies, which occupy an intermediate setting between pseudobulk analyses that collapse cells into biological samples and cell-level analyses that treat individual cells as independent observations. Accordingly, the mixed-effects framework accounts for correlation among cells arising from shared biological samples, tissue sections, or imaging runs while propagating uncertainty in transcript composition and global transcript abundance. This represents a simplifying assumption that these sources of correlation dominate the replicated study designs considered here. In study designs where the primary inferential target varies within a single tissue section, such as comparisons between neighboring anatomical regions or cellular neighborhoods, local spatial dependence may become a more important source of correlation. Because ALDEx interfaces with mixed-effects frameworks that support additional correlation structures, such as the spatial correlation classes available in nlme (Pinheiro et al., 2021), future implementations could jointly model spatial covariance alongside compositional uncertainty and biological replication when warranted by the study design. The reproducibility of both gene-level and pathway-level findings across independent MERFISH and CosMx datasets suggests that the current framework captures the dominant sources of uncertainty for replicated between-sample studies while remaining compatible with richer covariance models when warranted. As with all cell-level analyses, inference also remains dependent on adequate biological replication and the accuracy of upstream segmentation and cell-type annotation.

## Conclusions

In this work, we demonstrate that compositional uncertainty and variation in total transcript abundance represent important but underappreciated sources of variability in spatial transcriptomics. By explicitly modeling these components within a mixed-effects framework, ALDEx enables calibrated between-sample differential-expression analysis while preserving cell-level resolution. Across simulations and independent aging datasets, scale-aware modeling improved statistical calibration, enhanced reproducibility of aging-associated signals, and produced transcriptional programs that were more coherent across multiple complementary representations of biological aging. More broadly, our findings support treating uncertainty in global transcript abundance as an explicit component of inference rather than a nuisance to be removed through normalization, improving both reproducibility and biological interpretability across spatial transcriptomic studies.

## Methods

### Datasets used to illustrate scale and compositional uncertainty

To assess scale variability across targeted spatial transcriptomic assays, we analyzed MERFISH, Xenium, and CosMx mouse brain datasets spanning different panel sizes and platforms. Xenium comparisons were performed using publicly available mouse brain datasets distributed through the 10x Genomics Xenium data portal, including (i) Fresh Frozen Mouse Brain Replicates (Xenium Mouse Brain Gene Expression Panel), (ii) Fresh Frozen Mouse Brain Hemisphere (5K Mouse Pan Tissue and Pathways Panel), and (iii) Xenium In Situ Analysis of Alzheimer’s Disease Mouse Model Brain Coronal Sections (Mouse Brain Gene Expression Panel plus custom add-on panel). Dataset URLs are provided in Additional File 2: Table S1. Additionally, we used the anterior coronal left-hemisphere section from the SenNet CosMx mouse brain dataset (SNT638.MWRV.378; Bernlohr et al., 2026). To reduce biological confounding, cross-platform comparisons were restricted to young adult mouse brain datasets and anatomically similar coronal sections where available. Differences in observed transcript abundance therefore primarily reflect variation in panel composition, assay chemistry, and measurement platform.

Cells with zero counts were removed, and each dataset was filtered to exclude the lowest and highest 10% of cells by cell volume or cell area, where available. For panel-size comparisons, analyses were restricted to genes shared across the compared panels before recalculating per-cell total counts. For gene-level relative-expression comparisons, mean expression for each shared gene was normalized by the average count per detected gene within each cell, and log10-transformed relative expression values were compared across panels or platforms. Spatial transcript-depth structure was evaluated in matched MERFISH coronal sections by comparing total transcript counts across anatomical regions and by calculating mean absolute differences in total counts between neighboring cells as a function of spatial distance.

To illustrate the limited reproducibility of aging-associated DE signatures across studies, we compared published aging gene sets from three large mouse brain single-cell transcriptomic atlases: Tabula Muris Senis (2020), Ximerakis et al. (2019), and Jin et al. (2024). Comparisons were performed within matched broad cell types (e.g. oligodendrocytes, endothelial cells, and microglia). For the Tabula Muris Senis and Ximerakis datasets, aging-associated genes were defined using an adjusted p-value threshold of 0.01 together with a study-specific effect-size cutoff derived from the empirical effect-size distribution. For the Allen Brain Institute dataset, published aging-associated gene sets corresponding to matched cell subtypes were used directly. Gene-set overlap was visualized using Venn diagrams.

### ALDEx model specification

We formulate our DE framework using a Bayesian partially identified model that separates uncertainty arising from transcript composition and uncertainty in global transcript abundance. Let *Y*_*n*_ ∈ *N*^*D*^ denote the observed count vector for cell *n*, where *D* is the number of genes in the panel. We denote the corresponding latent transcript abundance vector *W*·_*n*_, where the dot indicates all genes in cell *n*.

#### Decomposition of absolute abundance

We formally represent log-abundance as the sum of compositional and scale components,

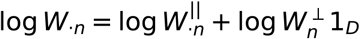

where 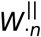 denotes the vector of relative transcript abundances across all genes in cell *n* and 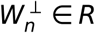 is a scalar latent variable representing transcriptome-wide abundance shared across all genes.

#### Measurement model for composition

Conditional on latent transcript composition *p*_*n*_, the observed transcript counts are modeled as

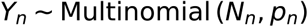

where 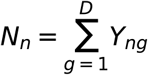 is the total number of transcripts detected in cell *n*, and *Y*_*ng*_ denotes the observed count for gene *g* in cell *n*.

A weakly informative symmetric Dirichlet prior is placed on *p*_*n*_,

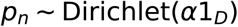

which yields the conjugate posterior,

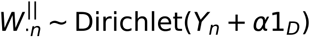

Monte Carlo samples from this posterior define realizations of the compositional component 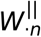, approximating uncertainty induced by finite sampling under compositional constraints.

#### Structured scale framework

Because global transcript abundance is not identifiable from observed counts alone, we represent uncertainty in the latent scale component using a Bayesian partially identified model.

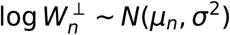

with expectation parameterized as

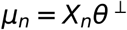

Here *X*_*n*_ denotes the row of the design matrix corresponding to cell *n, θ*^⊥^ ∈ *R*^*P*^ and contains the scale coefficients associated with the *P* predictors.

To propagate uncertainty in scale, for each Monte Carlo instance, scale coefficients are assigned a prior

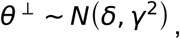

where *δ* specifies the prior mean and controls prior variability in transcriptome-wide scaling associated with predictors. In the absence of prior biological knowledge, *δ* = 0, yielding a zero-centered prior. Each sampled *θ*^⊥^ is mapped through the design matrix *X* to generate cell-specific realizations of 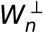.

#### Informed scale priors

When prior biological evidence suggests directional transcriptome-wide shifts in RNA abundance associated with specific predictors, the corresponding elements of are *δ* assigned non-zero values. For example, for the age predictors

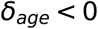

reflects prior evidence for reduced global RNA abundance with aging, while uncertainty in the magnitude of this effect is retained through the prior variance parameter *γ*^2^. Thus, the informed prior is a special case of the general scale prior above, differing only in the prior mean assigned to selected coefficients rather than introducing a separate statistical model.

Importantly, the scale component influences inference only through propagated uncertainty and does not directly determine gene-specific effect estimates.

#### Inference procedure

Inference is performed by jointly propagating uncertainty in compositional and scale components. For each Monte Carlo instance *s*, a compositional draw 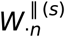 and a corresponding scale realization 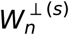 are combined to reconstruct transcript abundance according to

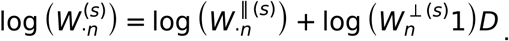

DE is then estimated using a mixed-effects model applied across Monte Carlo realizations,

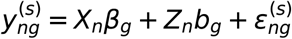

where 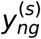 denotes the log-abundance for gene *g* in cell *n* under Monte Carlo sample *s, X*_*n*_ is the fixed effect design matrix, *Z*_*n*_ specifies the random effect structure and *b*_*g*_ represents gene-specific random effects.

Mixed-effects models are fit separately for each Monte Carlo realization. Gene-level statistics are aggregated to obtain posterior estimates of effect size and significance, and p-values are computed by combining test statistics across Monte Carlo samples using the ALDEx framework, followed by multiple testing corrections (e.g., Benjamini–Hochberg).

Mixed-effects models were fit using either the exact *lme4* engine (Bates et al., 2015) or the approximate *blmm* engine implemented in ALDEx3. The *lme4* engine performs full variance-component optimization independently for each Monte Carlo realization, providing exact inference but incurring substantial computational cost when the number of cells, features, or Monte Carlo samples is large.

The *blmm* engine accelerates inference by replacing repeated variance-component optimization across Monte Carlo draws with a feature-specific anchor fit followed by draw-specific local updates to covariance parameters, while retaining exact conditional estimation of fixed effects. This approximation substantially reduces runtime while targeting the same underlying mixed-effects model. The computational advantage of *blmm* is most pronounced for analyses with large numbers of Monte Carlo realizations, high-dimensional feature spaces, or computationally intensive random-effects structures.

For primary analyses, including the main MERFISH aging dataset and the Sun et al. subset, we used the exact *lme4* engine to ensure consistency with established inference. For computationally intensive secondary analyses, including large-scale simulations, cross-study comparisons, and high-dimensional subsets, we used the *blmm* engine to improve scalability. Results obtained using the approximate *blmm* engine were highly consistent with those from the exact *lme4* engine on representative subsets of the primary dataset (Additional File 4).

In practice, the *lme4* engine resulted in runtimes on the order of multiple days for large cell populations and complex random-effects structures, whereas the *blmm* approximation reduced runtime by over an order of magnitude, enabling large-scale simulation and cross-study analyses that would otherwise be computationally impractical. In settings with very large sample sizes and highly complex grouping structures, certain nuisance grouping variables were alternatively modeled as fixed effects to further improve computational tractability.

### Benchmarking under exchangeable nulls and controlled signal

We constructed a simulation framework to evaluate calibration and sensitivity of DE methods under realistic spatial transcriptomic measurement constraints. Beyond evaluating error control and sensitivity, these benchmarking experiments serve as an empirical diagnostic of identifiability under realistic spatial transcriptomic study designs. Simulations were based on a ST dataset measuring gene expression across brain sections from young and old mice. The dataset consists of six biological replicates (mice), each contributing two sections, yielding 12 samples organized into six imaging batches, with each batch containing a matched young and old section. Cells were assigned to annotated cell types and anatomical regions using the preprocessing and spatial annotation procedures described in the Application Dataset and Analysis Pipeline section. DE analyses operate at the cell level within cell type × region strata, with batch corresponding to imaging runs of paired tissue sections (Additional File 3).

#### Simulation design

To evaluate DE behavior under realistic study designs, we first generated exchangeable null datasets by permuting age labels within batches so that the association between age and expression was removed while maintaining the spatial, batch, and compositional structure of the data.

Because repeated refitting across all strata and simulation conditions is computationally intensive, simulations were performed within a single representative cell type × region stratum (astrocytes in the isocortex). To examine performance across study sizes, we constructed a grid of sample-size scenarios corresponding to N= 4, 8, 16, 32, 64, and 100 batches. For each scenario, batch units were sampled with replacement to generate datasets of the desired size (Additional File 3). Cells belonging to the selected batches were duplicated as needed to form the resampled dataset. Age labels were then permuted within each resampled batch instance to ensure that permutations did not mix independent duplicated copies.

Controlled gene-specific signal was then introduced on top of these exchangeable null datasets. For each simulation replicate, 20% of genes were selected as differentially expressed.

Differential effects were assigned asymmetrically, with approximately 80% downregulated and 20% upregulated in the Old group, reflecting the expectation that aging is associated with a general reduction in transcriptional output. Per-gene effect magnitudes were drawn uniformly from a realistic range on the log2 scale (approximately 0.5–4) (Additional File 3).

To ensure full reproducibility, each simulation replicate was defined by a simulation design table (Additional File 3) specifying independent random seeds for batch resampling, label permutation, differential gene selection, thinning, and multinomial resampling. Prior to model fitting, genes expressed in fewer than 20% of cells were excluded.

#### Thinning procedure

Gene-specific signal was introduced using binomial thinning as implemented in the seqgendiff framework (Gerard, 2020). The function thin_diff() was used to modify counts while preserving the original count structure and experimental design. Modified counts were generated as

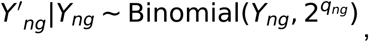

Where *Y*_*ng*_ and 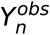 denote the observed and thinned counts, respectively, for gene *g* in cell *n*. The quantity *q*_*ng*_ encodes the linear predictor specifying the desired log2-fold change based on the design matrix and specified coefficients, with values shifted to ensure valid binomial probabilities. In our implementation, the permuted age labels define a binary design variable for signal injection, together with a corresponding gene-by-covariate coefficient vector specifying the injected log2 fold changes for DE genes (and zero for non-DE genes).

Thinning was applied after constructing each resampled dataset via batch-level up- or down-sampling and after restricted label permutation, so that the injected signal was evaluated under the same batch structure and exchangeability constraints used for calibration.

After signal injection by binomial thinning, we added a multinomial resampling step to better approximate the measurement process and restore the finite-budget compositional measurement structure of the observed data. Specifically, the thinned counts for each cell were treated as a latent abundance profile rather than as the final observed counts. For each cell,

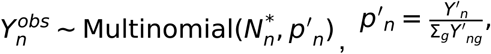

where 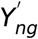 denotes the final observed count vector after multinomial resampling, *N* \**n* is the sampled total transcript counts for cell *n*, and 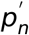 is the normalized abundance vector obtained from the thinned counts. The sampled totals *N*\**n* were drawn from the empirical distribution of per-cell total transcript counts within the corresponding batch.

Direct thinning alone modifies counts on an effectively absolute scale, without imposing the finite-count constraint that induces compositional coupling among genes. Multinomial resampling restores this constraint, producing observed counts that reflect both the injected signal and the stochastic measurement process.

#### ALDEx benchmarking specification

Within the benchmarking framework, ALDEx was fit separately for each simulated dataset using reduced Monte Carlo sampling (250 instances per stratum) to balance inferential stability with computational tractability across the full simulation grid. Because age labels were repeatedly permuted and cells duplicated during resampling, DE models were fit without cell-volume adjustment. Both the agnostic structured scale model and the informed structured scale formulation were evaluated to compare uncertainty propagation under alternative assumptions about global RNA shifts.

#### Other DE methods and benchmarking metrics

For each signal-injected replicate × sample-size scenario, simulated datasets were exported in interoperable count-matrix formats to enable benchmarking across DE frameworks implemented in both R and Python. MAST was applied in R at the cell level using age as the primary predictor under a hurdle model framework (Finak et al., 2015), whereas Wilcoxon rank-based testing was performed in Python using Scanpy’s rank_genes_groups implementation on the matched simulated count matrices (Wolf et al., 2018). A generalized linear mixed-effects negative binomial model was additionally implemented in R using the glmmTMB framework (Brooks et al., 2017) to model over dispersed count data while accounting for batch-level variability through random intercepts. Models were fit at the cell level using age as the primary fixed effect and total observed transcript counts as a log-offset term to account for differences in sequencing depth across cells. Gene-wise statistics from each framework were recorded and evaluated under known signal ground truth using empirical FDR, power, positive predictive value (PPV), and precision-oriented F-scores (Additional File 3). Metrics were summarized as a function of sample size across repeated simulation replicates.

### Spatial Transcriptome Data Collection and Analysis Pipeline

#### Primary MERFISH aging dataset overview

All animal care and experimental procedures were approved by the Penn State University Institutional Animal Care and Use Committee (IACUC). Spatial transcriptomic data were generated from C57BL/6J mice assigned to two age groups: young (n = 3; 2–3 months; 2 males, 1 female) and old (n = 3; 22 months; 2 males, 1 female). One young and one old mouse were processed together per cryosectioning session. From each mouse, two hemicoronal brain sections were collected: one from an anterior plane (bregma: ∼+0.98 mm) and one from a posterior plane (bregma: ∼−1.06 mm), yielding 12 sections total across all animals.

Each mouse was euthanized by cervical dislocation, immediately decapitated, and the brain was carefully dissected. The dissected brain was placed on ice while the procedure was repeated for the second mouse. Once both brains were extracted (∼5–8 minutes total), a distinguishing cut was made to facilitate sample identification after co-processing: the young brain was sectioned in a sagittal plane one section lateral to the midline, while the old brain was always cut at the midline. Dissection tools and blades were treated with RNaseZap (Invitrogen, AM9780) before and after each use to preserve RNA integrity. For tissue embedding, OCT compound (Tissue-Tek, 4853) was added to fill approximately one-fifth of a cryomold (Ted Pella, 27110). The young and old hemibrains were then carefully positioned on the OCT within the mold such that both hemispheres were in close apposition, approximating the appearance of an intact whole brain and the gross dorsal landmarks—olfactory bulb, cortex, and cerebellum— were visually aligned across both samples. The anterior–posterior (A/P) axis was noted on the cryomold before filling the remainder of the mold with OCT. The mold was rapidly frozen in 2-methylbutane pre-cooled with dry ice. Once fully solidified, the block was labeled and stored at −80°C until further use.

Coronal sections of 10 µm thickness were collected using a cryostat in the anterior-to-posterior direction, beginning at the olfactory bulb. As sectioning progressed, mediolateral and anteroposterior tilt was adjusted iteratively to ensure that the cutting plane through the young and old hemibrains produced sections at equivalent anatomical levels. Once the region of interest was approached, every 10th section was mounted on a glass slide and stained with DAPI. These DAPI-stained sections were examined to verify that key anatomical landmarks matched the target bregma coordinates and were concordant between the young and old hemibrains before proceeding to MERSCOPE slide collection. Following confirmation, 3–4 consecutive sections were mounted on MERSCOPE slides (Vizgen 2040000), ensuring that the entire tissue section fell within the imageable area. Approximately 10 sections collected immediately after the MERSCOPE slide series were pooled into an Eppendorf tube and stored at −80°C for RNA quality assessment. Prior to MERFISH runs, RNA integrity was evaluated by extracting RNA from these sections using the RNeasy Mini Kit (Qiagen, Cat. No. 74104) and determining RNA Integrity Numbers (RIN) on a 2100 Bioanalyzer using the RNA Nano chip (Agilent). Only samples with RIN ≥ 7 were advanced to MERFISH analysis. This workflow was completed first for the anterior target region and then repeated for the posterior target region.

Sections mounted on MERSCOPE slides were fixed in 4% paraformaldehyde (PFA) in 1× PBS for 15 minutes, washed three times with 1× PBS, and permeabilized in 70% ethanol at 4°C overnight. Subsequent sample preparation for MERSCOPE runs followed established protocols (Manjila et al., 2025; Moffitt and Zhuang, 2016; Moffitt et al., 2018); comprehensive details are available through Vizgen Resources and Vizgen Instrumentation documentation. A custom 500-gene panel was employed, encompassing cell-type markers and genes associated with cerebrovascular function, energy metabolism, and glycosylation pathways.

#### Spatial Transcriptomics Analysis Pipeline

ST preprocessing and analysis were performed using a Python framework developed for this project (*kimlabspatial*), which integrates Scanpy and Squidpy (Palla et al., 2022) functionality with customized modules for preprocessing, hierarchical cell-type identification, DE modeling, and visualization. The framework was developed to streamline ST workflows, including pooling datasets, bulk quality control, doublet detection, hierarchical clustering for cell typing, preparation of count matrices for downstream ALDEx modeling and postprocessing DE analysis results.

##### Registration to the Allen CCF

Brain sections were registered to the Allen Common Coordinate Framework (CCF v3, 2017 delineations; Q. Wang et al., 2020). Nuclear staining (DAPI) images were exported from the MERFISH Visualizer together with spatial metadata. Sections were aligned to the CCF using QuickNII (RRID:SCR_016854; Puchades et al., 2025) for linear alignment and estimation of the cutting plane, followed by nonlinear refinement using VisuAlign (RRID: SCR_017978; Gurdon et al., 2024). Registration accuracy was assessed using anatomical landmarks such as the corpus callosum, olfactory tubercle, and striatal structures, together with the spatial expression patterns of canonical marker genes (e.g., *Gad1, Slc17a7, Slc6a11*) to verify alignment of major anatomical boundaries. The final VisuAlign outputs were converted into 16-bit atlas label images using custom Python scripts, enabling assignment of each cell and transcript to a CCF brain region.

##### Cell Segmentation

Initial watershed-based segmentations produced by the MERFISH pipeline were refined using Cellpose-based segmentation (Stringer et al., 2021). Via the Vizgen Postprocessing Tool (VPT), custom Cellpose2 model (Pachitariu and Stringer, 2022) was trained using manually annotated cell boundaries to reduce misclassification of spherical imaging artifacts. Segmentation quality was evaluated on randomly selected image patches and compared to both the original watershed segmentation and the default Cellpose model. The custom Cellpose2 model achieved the highest accuracy (F1 ≈ 0.90) and was therefore used for downstream analyses.

##### Quality Control and Filtering

Preprocessing was performed using Scanpy-based workflows implemented within kimlabspatial. Datasets from all batches were pooled, and putative doublets were identified using Scrublet (Wolock et al., 2019). Cells were filtered based on segmentation volume, transcript counts, and gene detection thresholds to remove low-quality segmentations and imaging artifacts. Segmentation volumes were restricted to the 0.1–0.9 quantile range (48–3,583 μm^3^). Additional filters removed cells expressing fewer than 30 transcripts or fewer than 5 genes and genes detected in fewer than three cells. After filtering, the dataset contained cells with a median of 576 transcripts and 155 detected genes perf cell.

##### Cell Type Identification

Cell types were identified using a semi-automated hierarchical clustering strategy guided by curated marker genes. Marker sets were derived primarily from the Allen Brain Cell Atlas (Yao et al., 2023) and adapted to the MERFISH gene panel.

Clustering was performed iteratively using a marker-informed hierarchical framework. Initial clustering separated broad cell classes using standard Scanpy preprocessing (log transformation, total-count normalization, scaling, PCA, UMAP, and Leiden clustering). Candidate clustering solutions were generated through a grid search over PCA dimensionality, neighborhood size, and Leiden resolution parameters. Clustering performance was evaluated using a combined score incorporating silhouette score and a marker-purity metric quantifying agreement between clusters and expected marker expression patterns.

Top-scoring clustering solutions were manually reviewed and used as the starting point for hierarchical refinement. Subgroups of related clusters were subsequently reclustered using the same pipeline to resolve finer subtypes. Final cell type labels were assigned based on dominant marker expression within the hierarchical marker structure.

For neurons, a separate hierarchical marker tree was constructed using subclass and layer-specific markers derived from the Allen atlas. Although the primary aging analysis focused on non-neuronal populations, neuronal subclasses were also resolved using the same hierarchical framework to produce a consistent cell-type taxonomy across the dataset.

##### Differential Expression Data Preparation and Postprocessing

DE analyses in the primary dataset were performed using the ALDEx framework with mixed-effects models executed in R on a high-performance computing cluster. For each cell type × broad region stratum, raw count matrices and analysis metadata were exported using utilities within *kimlabspatial*. The primary aging analysis used the informed structured scale formulation with 2,000 Monte Carlo instances per stratum and mixed-effects models incorporating age and volume as fixed effects, and batch modeled as a random intercept (Additional File 4).

ALDEx outputs were postprocessed across multiple levels of biological organization, including cell type × region strata, cell types collapsed across regions, and broader summaries integrating matched vascular and immune-associated populations. Given the targeted nature of the MERFISH panel and the limited number of significant genes in many strata, functional interpretation was performed as a descriptive annotation summary rather than a formal enrichment analysis. GO and KEGG annotations associated with significant genes were aggregated across regions within each cell type and consolidated into broader functional modules. For each cell type, module prominence was computed by distributing each gene’s contribution across its associated annotations (normalized to sum to one per gene) and weighting each gene by the number of regions in which it was identified as significant. These weighted contributions were then aggregated within modules and normalized by the total contribution across all modules (Additional File 4).

### Cross-study reproducibility framework

To evaluate the reproducibility of scale-aware DE signals across independent spatial transcriptomic studies, we compared aging-associated effects from our primary dataset against two external aging-related imaging-based ST studies spanning different technologies, panel designs, section orientations, and age ranges.

#### Cross-study ALDEx specification

For all cross-study analyses, ALDEx was fit separately within matched cell type × broad region strata using 2,000 Monte Carlo instances per stratum and the informed structured scale formulation. Across all datasets, genes were retained only if expressed in at least 20% of cells within the stratum. Model formulas were matched as closely as possible to the available metadata in each study: the primary Sun comparison used binarized age and volume as fixed effects, and batch/cohort as a random effect (Additional File 5). Similarly, the SenNet Cosmx dataset comparison used age and segmented cell area as fixed effects, and paired experimental run as a random effect, closely matching the primary MERFISH dataset specification (Additional File 6).

#### Sun et al. coronal aging dataset

As a within platform reproducibility dataset, we used the MERFISH mouse brain aging study from Sun et al. (Nature, 2025), which profiled ∼2.3 million high-quality cells across 20 ages (3.4–34.5 months) in male C57BL/6JN mice using a 300-gene panel enriched for cell-type markers and aging-related pathways. To maximize comparability between datasets, cross-study analyses were restricted to the Sun coronal subset and an anatomically matched anterior-only batches of our primary dataset. Within the Sun dataset, we selected a subset of coronal samples spanning young (3.4–5.4 months) and old (19.8–24.6 months) to match the binary young-versus-old design of the primary aging dataset. Preprocessing steps were harmonized as closely as possible by applying equivalent cell-level quality filters and restricting analyses to shared non-neuronal populations. Cross-study comparisons were performed across matched broad regions, with primary emphasis on the isocortex, where anatomical correspondence between datasets was strongest (Additional File 5). In addition to binary cross-study comparisons, the full continuous-age Sun dataset was used for trajectory-consistency and functional-consistency analyses (Additional File 5).

#### SenNet Consortium CosMx sagittal aging dataset

To evaluate cross-platform reproducibility of aging-associated transcriptional programs, we reanalyzed a publicly available CosMx Spatial Molecular Imaging (SMI) mouse brain aging dataset generated by investigators at Mayo Clinic through the SenNet Consortium and described by Carver et al. (2026). The dataset consisted of 5-μm formalin-fixed, paraffin-embedded sagittal brain sections from four young (4 month) and four aged (24 month) female C57BL/6 mice profiled using the 950-plex CosMx Mouse Neuroscience panel supplemented with a custom 50-gene senescence-associated probe set. Imaging was performed at 120 nm/pixel resolution using the CosMx Spatial Molecular Imager platform with morphology-guided cellular segmentation and transcript assignment to polygonal cellular boundaries.

Raw transcript count matrices, metadata, and associated histological images were downloaded from the SenNet data portal. When these analyses were initiated, the dataset’s primary analysis had not yet been published; preprocessing, quality control, and broad cell-type annotation were therefore performed independently from the publicly available raw data. Metadata, tiled field-of-view organization, and sample pairing relationships were integrated to reconstruct the experimental design and anatomical organization of the dataset. Four paired young–old experimental runs were identified, each containing anterior and posterior sagittal regions of interest corresponding primarily to hippocampal/corticolimbic and cerebellar regions, respectively. Approximate anatomical planes and regional coverage were assessed using the Paxinos and Franklin Mouse Brain Atlas (Paxinos and Franklin, 2012).

Preprocessing and quality-control analyses were performed using Scanpy-based workflows implemented within *kimlabspatial*, paralleling the primary MERFISH analysis pipeline where possible. Raw CosMx transcript matrices were converted into AnnData format (Virshup et al., 2022). Cells were filtered based on transcript count and segmented cell area metrics, putative doublets were excluded prior to downstream analysis, and quality-control summaries were visually inspected across samples and anatomical regions.

Cell-type annotation was performed using complementary reference-mapping approaches based on both the Allen Institute’s MapMyCells framework (RRID:SCR_024672) and scANVI-based label transfer (Xu et al., 2021). Prior to mapping, gene identifiers were harmonized to Ensembl mouse gene IDs using MyGeneInfo-based symbol conversion (Wu et al., 2013). Because the sagittal CosMx regions of interest were anatomically restricted, non-overlapping taxonomy branches were excluded using the nodes_to_drop parameter. Multiple MapMyCells configurations were evaluated, including hierarchical versus flattened taxonomy assignment and alternative bootstrap settings. Final labels were defined at the class level for neuronal populations and subclass level for glial and vascular-associated populations (Astro-Epen, OPC-Oligo, and Vascular), reflecting the greater heterogeneity of non-neuronal classes (Additional File 2: Figure S3).

Consensus labels across MapMyCells configurations were computed using confidence-weighted agreement based on bootstrap support. scANVI predictions were subsequently used to adjudicate low-confidence assignments. Low-confidence assignments were retained when supported by concordant scANVI predictions or when both methods agreed at the broad neurotransmitter-class level. Remaining discordant ambiguous cells were excluded from downstream analyses.

DE analysis was performed using the ALDEx informed structured-scale model to maintain consistency with the primary MERFISH analysis. Analyses were performed separately for each retained cell type, including selected neuronal subclasses, using genes expressed in at least 20% of cells within the target population. Cell types represented by fewer than 1,000 cells across the dataset were excluded from analysis. Models included age and segmented cell area as fixed effects and paired experimental run as a random intercept. All models were fit using the *blmm* backend with 2,000 Monte Carlo instances per cell type (Additional File 6).

#### Cross-study concordance metrics

Reproducibility was quantified on the intersecting gene set for each matched comparison using complementary directional, ranking-based, and effect-size metrics. For the Sun dataset, comparisons were performed within matched cell type × region strata (Additional Files 4-5). For the CosMx dataset, comparisons were performed at the cell-type level owing to differences in anatomical organization and field-of-view structure. Directional reproducibility was evaluated using sign concordance, defined as the proportion of intersecting genes with matched effect directions between studies (Additional Files 4 and 6). Agreement among the strongest signals was assessed using CAT@K (concordance-at-the-top), defined as the fraction of overlap among the top K genes ranked by absolute effect size in each dataset, evaluated across a range of K values. Specifically, for each K, CAT@K was calculated as the number of shared genes among the top K ranked genes in both datasets divided by K. To directly compare reproducibility between analytical frameworks, differential CAT@K values were additionally computed as ΔCAT@K = CAT@K(ALDEx) − CAT@K(reference method), where positive values indicate greater overlap among top-ranked genes for the ALDEx framework. Effect-size concordance was summarized using Pearson and Spearman correlations of age-associated effect estimates across shared genes (Additional File 2: Tables S2-3).

#### Baseline comparison using rank-based pseudobulk analyses

In the original Sun et al. analysis, aging-associated genes were identified using Spearman correlation between age and pseudobulk expression across the full continuous age range. To provide a comparator framework for cross-study reproducibility analyses, we adapted this rank-based pseudobulk approach to a two-group setting and applied it to both the Sun and CosMx datasets (Additional File 4-6). Expression was normalized to a fixed total count, log-transformed, pseudobulked at the biological replicate level within each matched cell type, and compared between young and old groups using a Wilcoxon rank-sum test. Cross-study reproducibility for this baseline was evaluated analogously to ALDEx using the same sign concordance, CAT@K, and effect-size concordance metrics.

#### Directional consistency with raw transcript abundance

To assess whether inferred DE directions were consistent with observable changes in unnormalized measurements, we compared gene-level aging directions from ALDEx and rank-based pseudobulk analyses against corresponding shifts in raw transcript abundance and expression frequency within both external validation datasets.

For each gene within a matched cell type, we computed (i) the difference in mean raw transcript counts between old and young cells and (ii) the difference in the fraction of cells expressing the gene between old and young groups. Expression fraction was defined as the proportion of cells with nonzero transcript counts for the corresponding gene.

For each method, genes classified as increasing or decreasing with age were evaluated for directional agreement with these raw abundance metrics. Agreement rates were summarized as the proportion of inferred age-associated genes whose inferred direction matched the corresponding direction of change in raw counts or expression frequency. Analyses were performed separately within each validation dataset and summarized across matched cell types. This analysis was intended as a descriptive consistency assessment rather than a primary inferential benchmark, as raw transcript abundance does not account for compositional effects, differences in cellular RNA content, or uncertainty in global transcriptional scale.

#### Functional consistency analyses (Sun dataset)

To assess whether inferred aging-associated programs converged with independently derived biological signals, we compared continuous-age ALDEx and Spearman-based DE analyses against two published frameworks from Sun et al.: trajectory classifications based on lifespan expression dynamics and GO-BP enrichment results from the spatial aging clock analysis (Sun et al., 2025).

To extend our analysis to continuous age, ALDEx was fit on the full Sun coronal dataset using age as a continuous predictor (Additional File 5). Age was rescaled to the unit interval [0,1], with 0 corresponding to the youngest and 1 to the oldest sampled age. This scaled age variable was incorporated into both the mean (composition) and scale components of the model. The scale component was parameterized such that global transcript abundance varied linearly across the observed age range, preserving the interpretation used in the binary analysis in which older samples exhibit an expected reduction in total transcript abundance relative to younger samples. This formulation assumes an approximately linear relationship between age and both gene-specific and global transcript abundance effects.

For trajectory-consistency analysis, DE results were compared to trajectory classifications from the Sun dataset, where each gene was assigned to a trajectory cluster based on its cell type × region–specific expression profile across 20 ages (TableS7_GeneClassificationTrajectory in Sun et al.). Trajectory clusters were mapped to directional categories based on their qualitative temporal patterns. Clusters exhibiting monotonic or late-life increases (increasing gradual, increasing late) were classified as increasing, whereas clusters exhibiting monotonic or early-life decreases (decreasing gradual, decreasing early, midlife decrease) were classified as decreasing. Non-monotonic clusters (early peak, midlife peak, late peak, midlife trough) and genes lacking trajectory assignments were excluded from directional analyses. For both ALDEx and Spearman-based analyses, agreement between inferred gene-level direction and trajectory direction was evaluated using sign concordance and enrichment-based metrics.

In the original Sun et al. study, spatial aging clocks were constructed by spatially smoothing gene expression across same-cell-type neighborhood graphs before fitting cell-type-specific lasso regression models to predict biological age from spatial transcriptomic profiles. Genes with positive and negative clock coefficients were then analyzed separately for GO-BP enrichment using Fisher enrichment tests implemented through the topGO framework (Sun et al., 2025; Alexa and Rahnenfuhrer, 2026).

Genes were ranked by continuous-age ALDEx effect estimates separately within each cell type. GO-BP enrichment analysis was then performed by minimally modifying the published enrichment script from Sun et al (Additional File 5). Specifically, the original workflow based on Spearman correlation coefficients and confidence intervals was adapted to instead use ALDEx-derived continuous-age effect estimates and associated significance statistics while preserving the same topGO enrichment framework, ontology definitions, Fisher testing procedure, and term filtering criteria used in the original study. Published GO-BP enrichment results from both the Spearman-based analysis and the spatial aging clock analysis were used directly from the Sun et al. supplementary materials.

Functional consistency between methods was evaluated at the cell-type level using overlap-based metrics computed on enriched GO-BP term sets. To compare ALDEx- or Spearman-derived enrichments against the published clock-gene enrichments, we computed a three-way precision metric defined as the fraction of enriched terms from a given method that overlapped simultaneously with both the published Spearman-based enrichments and the published clock-gene enrichments (Additional File 2: Fig S2). Higher values therefore indicate stronger concentration within biological processes independently supported across all three analytical frameworks.

#### Functional annotation and module analysis (CosMx Dataset)

Functional interpretation was performed using a descriptive annotation-summary framework analogous to that applied in the primary MERFISH dataset. Given differences in gene-panel composition between platforms and the limited number of significant genes within several cell types, formal enrichment testing was not performed. Instead, GO-BP and KEGG pathway annotations associated with significant genes were retrieved using MyGeneInfo (Wu et al., 2013) and aggregated within each cell type (Supplementary Data S4). Recurrent annotations were manually consolidated into broader functional modules representing related biological programs.

To reduce bias from genes associated with large numbers of annotations, each gene’s contribution was distributed across its associated annotations such that contributions summed to one per gene. Annotation-level scores were computed by summing these fractional contributions across contributing genes, and module-level scores were obtained by aggregating annotation scores within each module. For cross-study functional comparisons, analyses were restricted to annotations shared between the MERFISH and CosMx datasets to facilitate comparison across platforms with differing gene-panel composition. Module-level concordance was evaluated using score- and rank-based comparisons across matched cell types, and representative annotation-level overlaps were examined to identify convergent biological programs supported by distinct underlying gene sets (Additional File 6).

#### Supplementary cross-study sensitivity analyses

Several additional sensitivity analyses were performed to assess the robustness of cross-study reproducibility conclusions. These included comparison of the adapted binary pseudobulk framework to the original continuous-age Spearman results, evaluation of anatomical-plane sensitivity within the primary dataset, resolution sensitivity using cell type–only summaries, and comparison between binary and continuous-age ALDEx results within the Sun dataset. Detailed results are provided in Additional Files 4-5.

## Declarations

**Acknowledgements**

We thank the authors of the Sun et al. aging MERFISH study for making their data and analysis resources publicly available. We acknowledge Carver et al., and the SenNet Consortium for generating and openly releasing the CosMx spatial transcriptomics datasets reanalyzed in this work. We thank members of the Kim laboratory for helpful discussions and feedback throughout the project. Additionally, we thank the high-performance computing (HPC) center, the Genome Sciences Core (RRID: SCR_021123) services with its instrument (MERSCOPE) used in this project at the Pennsylvania State University College of Medicine.

### Authors’ contributions

**Deniz Parmaksiz:** Conceptualization, Data curation, Formal analysis, Investigation, Methodology, Software, Visualization, Writing – original draft, Writing – review & editing.

**Steffy B. Manjila:** Data curation, Investigation, Resources, Writing – review & editing.

**Kyle C. McGovern:** Methodology, Software, Writing – review & editing.

**Donghui Shin:** Investigation, Writing – review & editing.

**Ingvild E. Bjerke:** Investigation, Writing – review & editing.

**Anirban Paul:** Resources, Writing – review & editing.

**Justin Silverman:** Conceptualization, Methodology, Supervision, Writing – review & editing.

**Yongsoo Kim:** Conceptualization, Funding acquisition, Project administration, Resources, Supervision, Writing – review & editing.

### Availability of data and materials

The processed primary MERFISH dataset and other data have been deposited in a Zenodo record. Source code and analysis scripts required to reproduce the analyses, including the spatial transcriptomics analysis pipeline and example workflows, are available through GitHub at https://github.com/denizparmaksiz/scale_aware_ST_DE.

Publicly available external datasets analyzed in this study were obtained from their original sources, including the Sun et al. MERFISH dataset, 10x Genomics Xenium public datasets, and SenNet CosMx datasets, including SNT249.TPCD.565. Dataset-specific citations, accession identifiers, URLs, and corresponding uses in the analyses are provided in the Methods and Additional File 2: Table S1.

### Ethics approval and consent to participate

Not applicable.

### Consent for publication

Not applicable.

## Supplementary Figure 1

**Figure S1.**
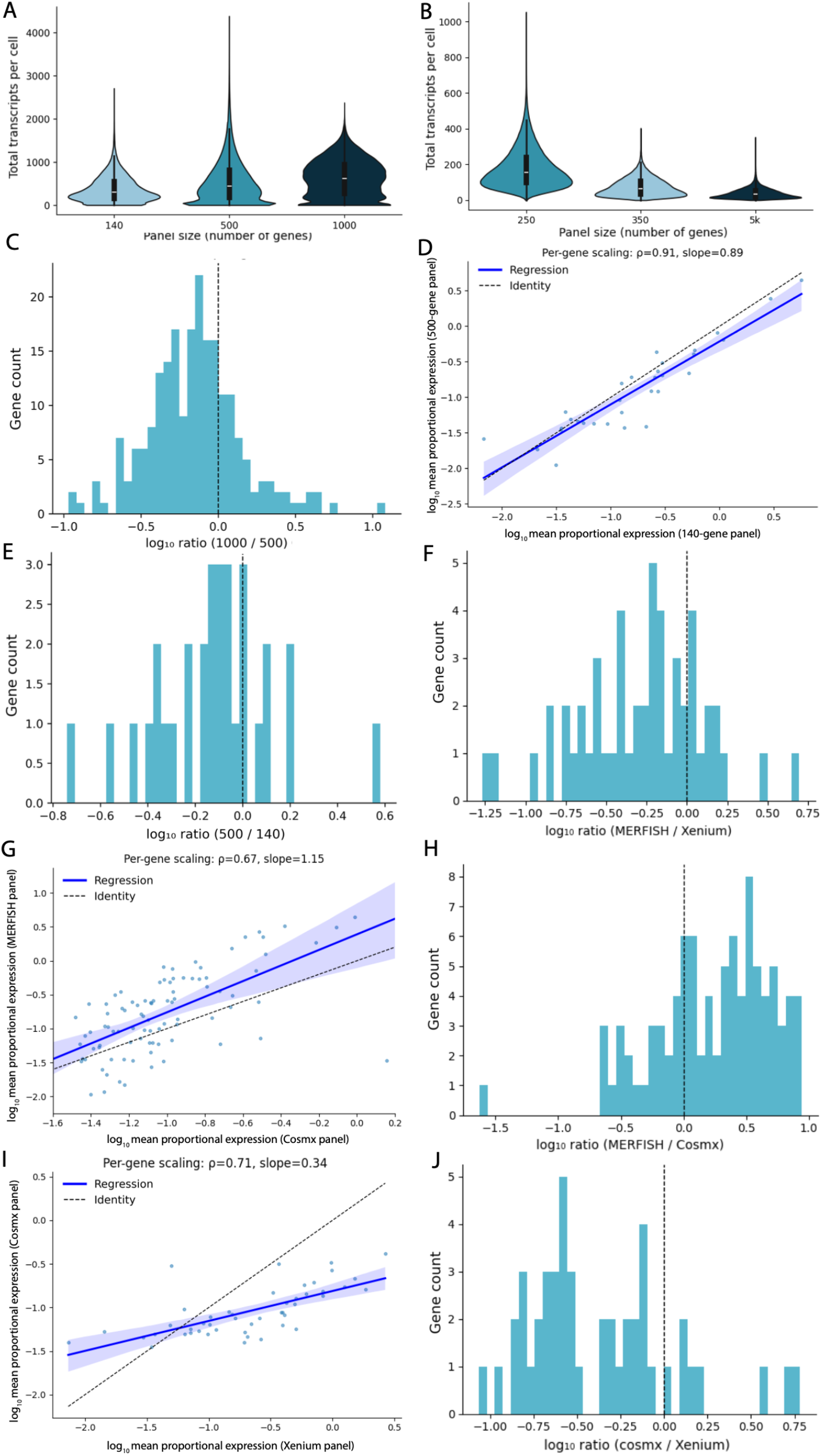
Additional evidence for scale uncertainty across ST platforms and panel designs. (a) Total transcripts detected per cell across MERFISH panels of increasing size (140-, 500-, and 1000-gene panels) using all genes in each panel (complementary to Fig. 1A). Transcript abundance increases with panel size but deviates from proportional scaling expectations. (b) Total transcripts detected per cell for genes shared across Xenium panels of different sizes. As observed for MERFISH (Fig. 1A), transcript counts for shared genes vary systematically with panel size despite restricting analysis to the same gene set. (c) Distribution of gene-specific shifts in proportional expression between the 500- and 1000-gene MERFISH panels. Histograms show log10 expression ratios for genes shared between panels, illustrating heterogeneous changes in gene detectability across panel designs. (d) Comparison of log_10_ mean proportional expression for genes shared between the 140- and 500-gene MERFISH panels. Deviations from the identity line indicate that proportional expression is not preserved across panel compositions. (e) Distribution of gene-specific shifts in proportional expression between the 140- and 500-gene MERFISH panels. Histograms show log10 expression ratios for genes shared between panels. (f) Distribution of gene-specific shifts in proportional expression between MERFISH and Xenium. Histograms show log10 expression ratios for genes shared across platforms, demonstrating heterogeneous platform-dependent differences in gene detectability. (g) Comparison of log_10_ mean proportional expression for genes shared between MERFISH and CosMx platforms. Deviations from the identity line indicate that proportional expression is not preserved across platforms. (h) Distribution of gene-specifishifts in proportional expression between MERFISH and CosMx. Histograms show log10 expression ratios for genes shared across platforms. (i) Comparison of log_10_ mean proportional expression for genes shared between Xenium and CosMx platforms. Deviations from the identity line indicate that proportional expression is not preserved across platforms. (j) Distribution of gene-specific shifts in prop between Xenium and CosMx. Histograms show log10 expression ratios for genes shared across platforms.

## Supplementary Figure 2

**Figure S2.**
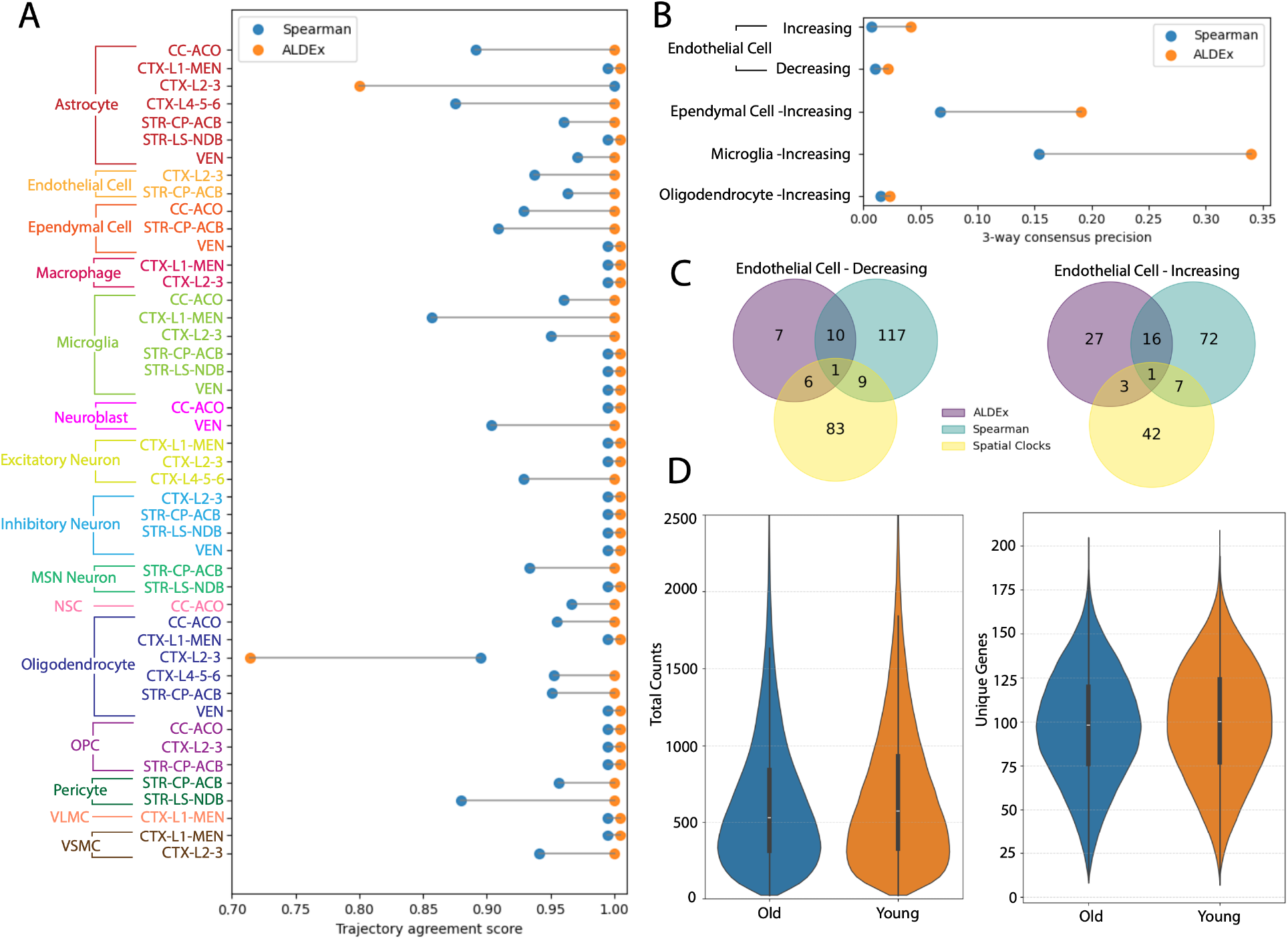
Supplementary validation analyses in the Sun et al. coronal aging dataset. (a) Agreement between inferred aging-associated expression changes and published lifespan trajectory classifications from the Sun et al. coronal aging dataset. For each matched cell type × subregion combination, points show the proportion of genes whose inferred direction of change agrees with the corresponding trajectory direction for continuous-age ALDEx and the published Spearman-based analysis. Connected points indicate matched comparisons. (b) Three-way consensus of age-associated GO-BP enrichments across ALDEx, published Spearman-based DE analysis, and spatial ageing clock genes. Connected points indicate matched cell type– direction comparisons, with higher values indicating greater agreement across independent analytical frameworks. (c) Example overlap of enriched GO-BP terms among ALDEx, published Spearman-based DE analysis, and spatial ageing clock genes for decreasing (left) and increasing (right) endothelial-cell ageing programs in the Sun coronal dataset. Although ALDEx identified fewer enriched GO-BP terms overall, a large proportion were shared with both alternative approaches, indicating enrichment for consensus ageing-associated biological programs. (d) Distribution of total transcript counts (left) and detected genes per cell (right) in matched young (3.4–5.4 months) and old (30.9–34.5 months) cohorts from the Sun coronal dataset. Age groups were selected from samples paired within experimental slides to minimize technical confounding. Old cells exhibited modest reductions in both total transcript abundance and the number of detected genes per cell relative to young cells.

## Supplementary Figure 3

**Figure S3.**
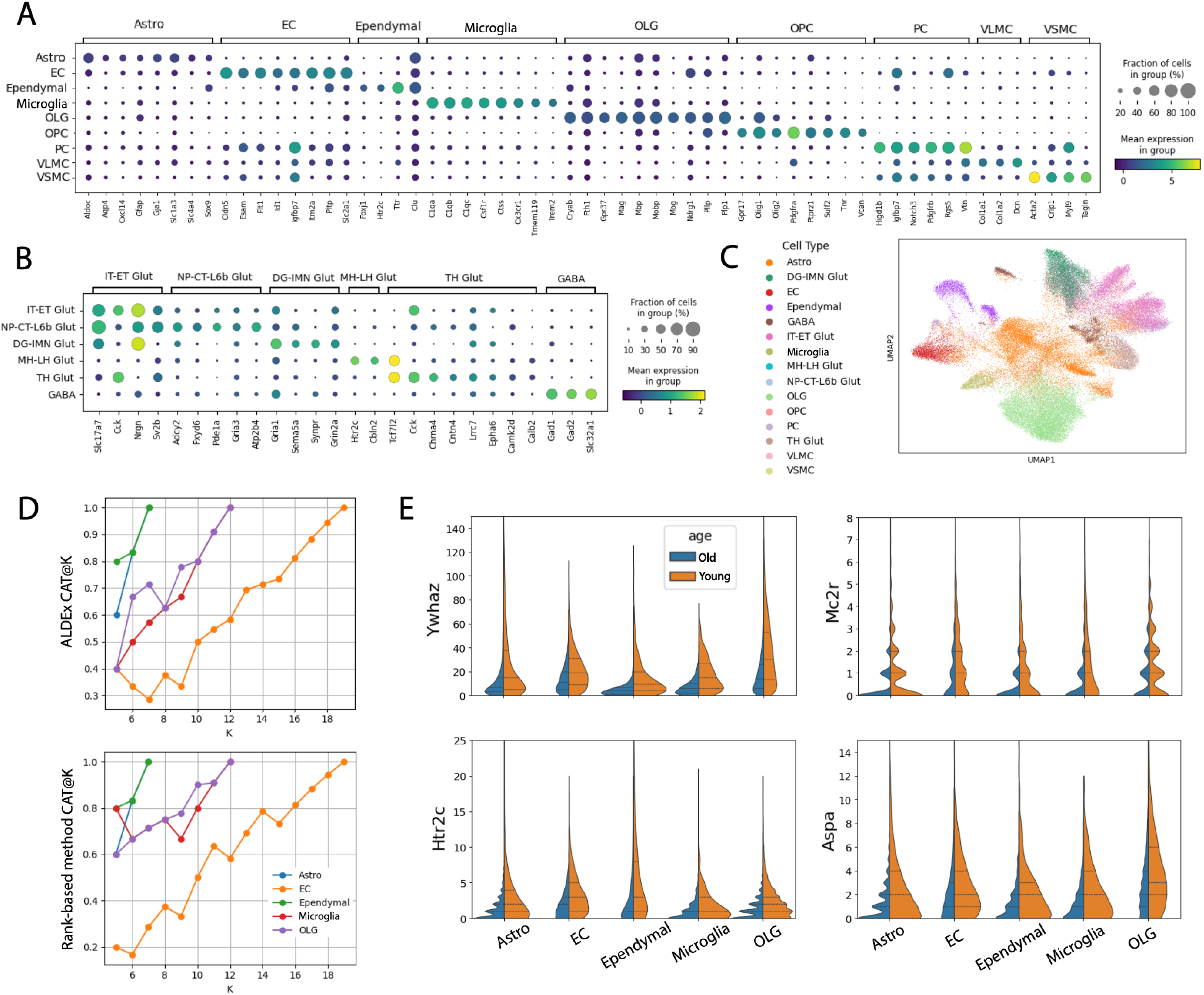
Cell-type annotation and cross-study reproducibility analyses for the CosMx SenNet aging dataset. (a) Dot plot of non-neuronal cell-type assignments in the CosMx SenNet aging dataset. Expression of representative marker genes is shown across broad glial, vascular, and immune populations following consensus reference mapping using MapMyCells and scANVI. (b) Dot plot of neuronal cell-class assignments in the CosMx SenNet aging dataset. Representative marker genes are shown for major excitatory and inhibitory neuronal classes following consensus reference mapping. (c) UMAP visualization of CosMx cells colored by final consensus cell-type assignments. Labels were derived using confidence-weighted integration of MapMyCells and scANVI predictions following quality control and filtering.(d) CAT@K analysis comparing overlap among top-ranked aging-associated genes between the primary MERFISH and CosMx datasets across a range of K values. Results are shown separately for ALDEx and the adapted rank-based pseudobulk framework, with higher values indicating greater agreement among highly ranked aging-associated genes across studies. (e) Additional examples of aging-associated genes identified in the independent CosMx SenNet aging dataset. Violin plots show raw transcript abundance distributions in young and aged cells for representative genes exhibiting consistent decreases or increases across multiple non-neuronal populations. Corresponding examples shown in Fig. 4E highlight recurrent age-associated transcriptional changes supported by shifts in observed transcript abundance.

## References

1. Aitchison J. The Statistical Analysis of Compositional Data. Chapman & Hall; 1986.

2. Alexa A, Rahnenführer J. topGO: Enrichment Analysis for Gene Ontology. 2026; doi:10.18129/B9.bioc.topGO. R package version 2.64.0, https://bioconductor.org/packages/topGO.

3. Allen Institute for Brain Science, 2023. MapMyCells, RRID:SCR_024672. Available at: < https://brain-map.org/bkp/analyze/mapmycells

4. Atta L, Clifton K, Anant M, Aihara G, Fan J, et al. Gene count normalization in single-cell imaging-based spatially resolved transcriptomics. Genome Biology. 2024;25:153. doi:10.1186/s13059-024-03303-w.

5. Bates, Douglas, et al. “Fitting Linear Mixed-Effects Models Using Lme4”. Journal of Statistical Software, vol. 67, no. 1, Oct. 2015, pp. 1–48, doi:10.18637/jss.v067.i01.

6. Bernlohr D, Niedernhofer L, DuFresne-To M, et al. CosMx Transcriptomics data from the brain of a C57BL/6 female mouse. Published online 2026. doi:10.60586/SNT638.MWRV.378

7. Bhuva DD, Tan CW, Salim A, Marceaux C, Pickering MA, Chen J, et al. Library size confounds biology in spatial transcriptomics data. Genome Biology. 2024;25:99. doi:10.1186/s13059-024-03241-7.

8. Bolaños JP, Almeida A. Signaling roles for astrocytic lipid metabolism in brain function. EMBO Rep. 2026 Feb;27(3):573–580. doi: 10.1038/s44319-025-00683-3. Epub 2026 Jan 3. PMID: 41484383; PMCID: PMC12894840.

9. Braunstein KE, Eschbach J, Ròna-Vörös K, Soylu R, Mikrouli E, Larmet Y, René F, Gonzalez De Aguilar JL, Loeffler JP, Müller HP, Bucher S, Kaulisch T, Niessen HG, Tillmanns J, Fischer K, Schwalenstöcker B, Kassubek J, Pichler B, Stiller D, Petersen A, Ludolph AC, Dupuis L. A point mutation in the dynein heavy chain gene leads to striatal atrophy and compromises neurite outgrowth of striatal neurons. Hum Mol Genet. 2010 Nov 15;19(22):4385–98. doi: 10.1093/hmg/ddq361. Epub 2010 Aug 31. PMID: 20807776; PMCID: PMC3298848.

10. Brooks ME, Kristensen K, van Benthem KJ, Magnusson A, Berg CW, Nielsen A, Skaug HJ, Maechler M, Bolker BM (2017). “glmmTMB Balances Speed and Flexibility Among Packages for Zero-inflated Generalized Linear Mixed Modeling.” The R Journal, 9(2), 378–400. doi:10.32614/RJ-2017-066.

11. Cable DM, Murray E, Shanmugam V, Zhang S, Zou LS, Diao M, et al. Cell type-specific inference of differential expression in spatial transcriptomics. Nature Methods. 2022;19:1076–1087.

12. Cao J, Li C, Cui Z, Deng S, Lei T, Liu W, Yang H, Chen P. Spatial Transcriptomics: A Powerful Tool in Disease Understanding and Drug Discovery. Theranostics. 2024 May 11;14(7):2946–2968. doi: 10.7150/thno.95908. PMID: 38773973; PMCID: PMC11103497.

13. Carver CM, Gomez PT, Rodriguez SL, et al. Spatial mapping and senolytic targeting of senescent and disease-associated microglia in aged mouse brain white matter. Nature Aging. 2026;6:1417–1436. doi:10.1038/s43587-026-01154-7.

14. Cheng Y, Cawley NX, Loh YP. Carboxypeptidase E/NFα1: a new neurotrophic factor against oxidative stress-induced apoptotic cell death mediated by ERK and PI3-K/AKT pathways. PLoS One. 2013 Aug 15;8(8):e71578. doi: 10.1371/journal.pone.0071578. PMID: 23977080; PMCID: PMC3744492.

15. Cui X, Liu S, Song H, Xu J, Sun Y. Single-cell and spatial transcriptomic analyses revealing tumor microenvironment remodeling after neoadjuvant chemoimmunotherapy in non-small cell lung cancer. Mol Cancer. 2025 Apr 9;24(1):111. doi: 10.1186/s12943-025-02287-w. PMID: 40205583; PMCID: PMC11980172.

16. Cunnane SC, Schneider JA, Tangney C, Tremblay-Mercier J, Fortier M, Bennett DA, Morris MC. Plasma and brain fatty acid profiles in mild cognitive impairment and Alzheimer’s disease. J Alzheimers Dis. 2012;29(3):691–7. doi: 10.3233/JAD-2012-110629. PMID: 22466064; PMCID: PMC3409580.

17. Elmarasi M, Elmakaty I, Elsayed B, Elsayed A, Zein JA, Boudaka A, Eid AH. Phenotypic switching of vascular smooth muscle cells in atherosclerosis, hypertension, and aortic dissection. J Cell Physiol. 2024 Apr;239(4):e31200. doi: 10.1002/jcp.31200. Epub 2024 Jan 30. PMID: 38291732.

18. Fan FC, D. Y, Zheng WH, Loh YP, Cheng Y. Carboxypeptidase E conditional knockout mice exhibit learning and memory deficits and neurodegeneration. Transl Psychiatry. 2023 Apr 26;13(1):135. doi: 10.1038/s41398-023-02429-y. PMID: 37100779; PMCID: PMC10133319.

19. Finak G, McDavid A, Yajima M, Deng J, Gersuk V, Shalek AK, Slichter CK, Miller HW, McElrath MJ, Prlic M, Linsley PS, Gottardo R. MAST: a flexible statistical framework for assessing transcriptional changes and characterizing heterogeneity in single-cell RNA sequencing data. Genome Biol. 2015 Dec 10;16:278. doi: 10.1186/s13059-015-0844-5. PMID: 26653891; PMCID: PMC4676162.

20. Gao P, Gao P, Choi M, Chegireddy K, Slivano O, Zhao J, Zhang W, Long X. Transcriptome analysis of mouse aortae reveals multiple novel pathways regulated by aging. Aging (Albany NY). 2020; 12:15603–15623. 10.18632/aging.103652

21. Gerard D. Data-based RNA-seq simulations by binomial thinning. BMC Bioinformatics. 2020 May 24;21(1):206. doi: 10.1186/s12859-020-3450-9. PMID: 32448189; PMCID: PMC7245910.

22. Grases D, Porta-Pardo E. A practical guide to spatial transcriptomics: lessons from over 1000 samples. Trends Biotechnol. 2026 May;44(5):1230–1242. doi: 10.1016/j.tibtech.2025.08.020. Epub 2025 Sep 19. PMID: 40975650.

23. Gurdon B, Yates SC, Csucs G, Groeneboom NE, Hadad N, Telpoukhovskaia M, Ouellette A, Ouellette T, O’Connell KMS, Singh S, Murdy TJ, Merchant E, Bjerke I, Kleven H, Schlegel U, Leergaard TB, Puchades MA, Bjaalie JG, Kaczorowski CC. Detecting the effect of genetic diversity on brain composition in an Alzheimer’s disease mouse model. Commun Biol. 2024 May 20;7(1):605. doi: 10.1038/s42003-024-06242-1. PMID: 38769398; PMCID: PMC11106287.

24. Gyenis A, Chang J, Demmers JJPG, Bruens ST, Barnhoorn S, Brandt RMC, Baar MP, Raseta M, Derks KWJ, Hoeijmakers JHJ, Pothof J. Genome-wide RNA polymerase stalling shapes the transcriptome during aging. Nat Genet. 2023 Feb;55(2):268–279. doi: 10.1038/s41588-022-01279-6. Epub 2023 Jan 19. PMID: 36658433; PMCID: PMC9925383.

25. Hafemeister C, Satija R. Normalization and variance stabilization of single-cell RNA-seq data using regularized negative binomial regression. Genome Biol. 2019 Dec 23;20(1):296. doi: 10.1186/s13059-019-1874-1. PMID: 31870423; PMCID: PMC6927181.

26. Jin K, Siletti K, Kleshchevnikov V, et al. Brain-wide cell-type-specific transcriptomic signatures of healthy ageing in mice. Nature. 2024. doi:10.1038/s41586-024-08350-8.

27. Kalantari-Dehaghi M, Ghohabi-Esfahani N, Emadi-Baygi M. From bulk RNA sequencing to spatial transcriptomics: a comparative review of differential gene expression analysis methods. Hum Genomics. 2025 Dec 6;20(1):9. doi: 10.1186/s40246-025-00884-w. PMID: 41353326; PMCID: PMC12797438.

28. Kiviaho A, Eerola SK, Kallio HML, Andersen MK, Hoikka M, Tiihonen AM, Salonen I, Spotbeen X, Giesen A, Parker CTA, Taavitsainen S, Hantula O, Marttinen M, Hermelo I, Ismail M, Midtbust E, Wess M, Devlies W, Sharma A, Krossa S, Häkkinen T, Afyounian E, Vandereyken K, Kint S, Kesseli J, Tolonen T, Tammela TLJ, Viset T, Størkersen Ø, Giskeødegård GF, Rye MB, Murtola T, Erickson A, Latonen L, Bova GS, Mills IG, Joniau S, Swinnen JV, Voet T, Mirtti T, Attard G, Claessens F, Visakorpi T, Rautajoki KJ, Tessem MB, Urbanucci A, Nykter M. Single cell and spatial transcriptomics highlight the interaction of club-like cells with immunosuppressive myeloid cells in prostate cancer. Nat Commun. 2024 Nov 16;15(1):9949. doi: 10.1038/s41467-024-54364-1. PMID: 39550375; PMCID: PMC11569175.

29. Kukreja B, Jeon S, Cao W, Rusu B, Harrison CF, Ghazisaeidi S, Tahmasian N, Feng MY, Chan R, Loan A, Johnston WB, Padhy S, Rakoff-Nahoum S, Wang J, Yim YS, Kalish BT. Spatial transcriptomics of the developing mouse brain immune landscape reveals effects of maternal immune activation and microbiome depletion. Nat Neurosci. 2026 Mar;29(3):732–745. doi: 10.1038/s41593-025-02162-3. Epub 2026 Jan 6. PMID: 41495254.

30. Li B, Tang Z, Budhkar A, Liu X, Zhang T, Yang B, Su J, Song Q. SpaIM: single-cell spatial transcriptomics imputation via style transfer. Nat Commun. 2025 Aug 23;16(1):7861. doi: 10.1038/s41467-025-63185-9. PMID: 40849313; PMCID: PMC12375071.

31. Li C, Chan T-F, Yang C, Lin Z. stVAE deconvolves cell-type composition in large-scale cellular resolution spatial transcriptomics. Bioinformatics. 2023;39(10):btad642. doi:10.1093/bioinformatics/btad642.

32. Li M, Wang H, Tang Z, Yang S, Lin L. Dysregulated Lipid Metabolism and Neurovascular Unit Dysfunction: Novel Mechanisms Linking Alzheimer’s Disease and Vascular Dementia. Aging Dis. 2026 Feb 13. doi: 10.14336/AD.2025.1464. Epub ahead of print. PMID: 41701875.

33. Li S, Chow LH, Pickering JG. Cell surface-bound collagenase-1 and focal substrate degradation stimulate the rear release of motile vascular smooth muscle cells. J. Biol. Chem. 2000;275:35384–35392. doi: 10.1074/jbc.M005139200.

34. Li S, Agudelo Garcia PA, Aliferis C, Becich MJ, Calyeca J, Cosgrove BD, Elisseeff J, Farzad N, Fertig EJ, Glass C, Gu L, Hu Q, Ji Z, Königshoff M, LeBrasseur NK, Li D, Ma A, Ma Q, Menon V, Mitchell JT, Mora AL, Nagaraj S, Nelson AC, Niedernhofer LJ, Rojas M, Taha HB, Wang J, Wang S, Wu PH, Xie J, Xu M, Yu M, Zhang X, Zhao Y, Adams PD, Aguayo-Mazzucato C, Baker DJ, Benz C, Bernlohr DA, Bueno M, Chen J, Childs BG, Chuang JH, Chung D, Dileepan M, Ding L, Dong M, Duncan F, Enninful A, Flynn WF, Franco AC, Furman D, Garovic V, Halene S, Herman AB, Hertzel AV, Iwasaki K, Jeon H, Kang JW, Karmakar S, Kirkland JL, Korstanje R, Kummerfeld E, Lee JH, Liu Y, Lu Y, Lugo-Martinez J, Martini H, Melov S, Musi N, Passos JF, Peters ST, Rahman I, Ramasamy R, Rindone AN, Robbins PD, Robson P, Rodriguez-Lopez J, Rosas L, Rosenthal N, Schafer MJ, Schilling B, Schmidt EL, Schneider K, Sengupta K, Shu J, So PTC, Sun L, Tchkonia T, Teneche MG, Vanegas N, Wang J, Xie J, Yin S, Zhang K, Zhu Q, Fan R; SenNet Consortium. Advancing biological understanding of cellular senescence with computational multiomics. Nat Genet. 2025 Oct;57(10):2381–2394. doi: 10.1038/s41588-025-02314-y. Epub 2025 Sep 15. PMID: 40954249; PMCID: PMC12916282.

35. Liu H, Jiang D, Yao F, Li T, Zhou B, Zhao S, Yang K, Feng H, Shen J, Tang J, Wang S, Zhang YX, Wang Y, Li Q, Zhao Y, Guo C, Tang TS. Restoring carboxypeptidase E rescues BDNF maturation and neurogenesis in aged brains. Life Med. 2023 Apr 11;2(2):lnad015. doi: 10.1093/lifemedi/lnad015. PMID: 39872114; PMCID: PMC11749474.

36. López-Otín C, Blasco MA, Partridge L, Serrano M, Kroemer G. Hallmarks of aging: An expanding universe. Cell. 2023 Jan 19;186(2):243–278. doi: 10.1016/j.cell.2022.11.001. Epub 2023 Jan 3. PMID: 36599349.

37. Makihara N, Arimura K, Ago T, Tachibana M, Nishimura A, Nakamura K, Matsuo R, Wakisaka Y, Kuroda J, Sugimori H, Kamouchi M, Kitazono T. Involvement of platelet-derived growth factor receptor β in fibrosis through extracellular matrix protein production after ischemic stroke. Exp Neurol. 2015 Feb;264:127–34. doi: 10.1016/j.expneurol.2014.12.007. Epub 2014 Dec 13. PMID: 25510317.

38. Manjila SB, Son S, Parmaksiz D, Kline H, Betty R, Wu Y-T, Pi H-J, Shin D, Liwang JK, Kronman FN, Bjerke IE, McGovern KC, Silverman JD, Paul A, Kim Y. Brain-wide connectivity and novelty response of the dorsal endopiriform nucleus in mice. Cell Reports. 2025;44(7):115827. doi:10.1016/j.celrep.2025.115827.

39. Martin N, Olsen P, Quon J, Campos J, et al. MerQuaCo: a computational tool for quality control in image-based spatial transcriptomics. eLife. 2025; 14:RP105149 10.7554/eLife.105149.1

40. Mason K, Sathe A, Hess PR, Rong J, Wu CY, Furth E, et al. Niche-DE: niche-differential gene expression analysis in spatial transcriptomics data identifies context-dependent cell–cell interactions. Genome Biology. 2024;25:14.

41. Massagué J, Sheppard D. TGF-β signaling in health and disease. Cell. 2023 Sep 14;186(19):4007–4037. doi: 10.1016/j.cell.2023.07.036. PMID: 37714133; PMCID: PMC10772989.

42. McGovern KC, Silverman JD. Scale reliant mixed effects models enhance microbiome data analysis. Microbiome. 2026. doi:10.1186/s40168-026-02377-x

43. Melo Dos Santos LS, Trombetta-Lima M, Eggen B, Demaria M. Cellular senescence in brain aging and neurodegeneration. Ageing Res Rev. 2024 Jan;93:102141. doi: 10.1016/j.arr.2023.102141. Epub 2023 Nov 27. PMID: 38030088.

44. Moffitt JR, Bambah-Mukku D, Eichhorn SW, Vaughn E, Shekhar K, Perez JD, Rubinstein ND, Hao J, Regev A, Dulac C, Zhuang X. Molecular, spatial, and functional single-cell profiling of the hypothalamic preoptic region. Science. 2018 Nov 16;362(6416):eaau5324. doi: 10.1126/science.aau5324. Epub 2018 Nov 1. PMID: 30385464; PMCID: PMC6482113.

45. Moffitt JR, Zhuang X. RNA Imaging with Multiplexed Error-Robust Fluorescence In Situ Hybridization (MERFISH). Methods Enzymol. 2016;572:1–49. doi: 10.1016/bs.mie.2016.03.020. Epub 2016 Apr 27. PMID: 27241748; PMCID: PMC5023431.

46. Mons N, Segu L, Nogues X, Buhot MC. Effects of age and spatial learning on adenylyl cyclase mRNA expression in the mouse hippocampus. Neurobiol Aging. 2004 Sep;25(8):1095–106. doi: 10.1016/j.neurobiolaging.2003.10.014. PMID: 15212834.

47. Moore SA, Yoder E, Murphy S, Dutton GR, Spector AA. Astrocytes, not neurons, produce docosahexaenoic acid (22:6 omega-3) and arachidonic acid (20:4 omega-6). J Neurochem. 1991 Feb;56(2):518–24. doi: 10.1111/j.1471-4159.1991.tb08180.x. PMID: 1824862.

48. Nakamura MT, Nara TY. Structure, function, and dietary regulation of delta6, delta5, and delta9 desaturases. Annu Rev Nutr. 2004;24:345–76. doi: 10.1146/annurev.nutr.24.121803.063211.

49. Nguyen LN, Ma D, Shui G, Wong P, Cazenave-Gassiot A, Zhang X, Wenk MR, Goh EL, Silver DL. Mfsd2a is a transporter for the essential omega-3 fatty acid docosahexaenoic acid. Nature. 2014 May 22;509(7501):503–6. doi: 10.1038/nature13241. Epub 2014 May 14. PMID: 24828044.

50. Nixon MP, Gloor GB, Silverman JD. Incorporating scale uncertainty in microbiome and gene expression analysis as an extension of normalization. Genome Biol. 2025 May 22;26(1):139. doi: 10.1186/s13059-025-03609-3. PMID: 40405262; PMCID: PMC12100815.

51. Ospina OE, Soupir AC, Manjarres-Betancur R, Gonzalez-Calderon G, Yu X, Fridley BL. Differential gene expression analysis of spatial transcriptomic experiments using spatial mixed models. Sci Rep. 2024 May 14;14(1):10967. doi: 10.1038/s41598-024-61758-0. PMID: 38744956; PMCID: PMC11094014.

52. Pachitariu M, Stringer C. Cellpose 2.0: how to train your own model. Nat Methods. 2022 Dec;19(12):1634–1641. doi: 10.1038/s41592-022-01663-4. Epub 2022 Nov 7. PMID: 36344832; PMCID: PMC9718665.

53. Palla G, Spitzer H, Klein M, Fischer D, Schaar AC, Kuemmerle LB, Rybakov S, Ibarra IL, Holmberg O, Virshup I, Lotfollahi M, Richter S, Theis FJ. Squidpy: a scalable framework for spatial omics analysis. Nat Methods. 2022 Feb;19(2):171–178. doi: 10.1038/s41592-021-01358-2. Epub 2022 Jan 31. PMID: 35102346; PMCID: PMC8828470.

54. Panov A, Orynbayeva Z, Vavilin V, Lyakhovich V. Fatty acids in energy metabolism of the central nervous system. Biomed Res Int. 2014;2014:472459. doi: 10.1155/2014/472459. Epub 2014 May 4. PMID: 24883315; PMCID: PMC4026875.

55. Park HE, Jo SH, Lee RH, Macks CP, Ku T, Park J, Lee CW, Hur JK, Sohn CH. Spatial Transcriptomics: Technical Aspects of Recent Developments and Their Applications in Neuroscience and Cancer Research. Adv Sci (Weinh). 2023 Jun;10(16):e2206939. doi: 10.1002/advs.202206939. Epub 2023 Apr 7. PMID: 37026425; PMCID: PMC10238226.

56. Paxinos G, Franklin KBJ. Paxinos and Franklin’s the mouse brain in stereotaxic coordinates. 4th ed. Academic Press, Cambridge, Massachusetts; 2012.

57. Pinheiro J, Bates D, DebRoy S, Sarkar D, R Core Team. nlme: Linear and Nonlinear Mixed Effects Models [Internet]. 2021. Available from: https://CRAN.R-project.org/package=nlme

58. Piwecka M, Rajewsky N, Rybak-Wolf A. Single-cell and spatial transcriptomics: deciphering brain complexity in health and disease. Nat Rev Neurol. 2023 Jun;19(6):346–362. doi: 10.1038/s41582-023-00809-y. Epub 2023 May 17. PMID: 37198436; PMCID: PMC10191412.

59. Plummer JT, Dezem FS, Cook DP, Park J, Zhang L, Liu Y, Marção M, DuBose H, Wani A, Wise K, Roach M, Harvey K, Wang T, Jensen KB, Morosini N, De Gregorio R, Alonso A, Houlihan SL, Schwartz RE, Hissong E, Snopkowski C, Wrana JL, Ryan N, Butler LM, Church G, Swarbrick A, Mason CE, Martelotto LG. Standardized metrics for assessment and reproducibility of imaging-based spatial transcriptomics datasets. Nat Biotechnol. 2025 Dec 3. doi: 10.1038/s41587-025-02811-9. Epub ahead of print. PMID: 41339526.

60. Puchades MA, Yates SC, Csucs G, Carey H, Balkir A, Leergaard TB and Bjaalie JG (2025) Software and pipelines for registration and analyses of rodent brain image data in reference atlas space. Front. Neuroinform. 19:1629388. doi: 10.3389/fninf.2025.1629388

61. Salim A, Bhuva DD, Chen C, Tan CW, Yang P, Davis MJ, et al. SpaNorm: spatially aware normalization for spatial transcriptomics data. Genome Biology. 2025;26:109. doi:10.1186/s13059-025-03565-y.

62. Salim A, Molania R, Wang J, De Livera A, Thijssen R, Speed TP. RUV-III-NB: normalization of single cell RNA-seq data. Nucleic Acids Research. 2022;50(16):e96. doi:10.1093/nar/gkac486.

63. Sarwar A, Rue M, French L, Cross H, Choi S, Chen X, Gillis J. Cross-expression meta-analysis of mouse brain slices reveals coordinated gene expression across spatially adjacent cells. Genome Biol. 2025 Oct 29;26(1):373. doi: 10.1186/s13059-025-03747-8. PMID: 41163012; PMCID: PMC12570847.

64. Shen Q, Dong K, Zhang S, Zhang S. High-precision cell-type mapping and annotation of single-cell spatial transcriptomics with STAMapper. Genome Biol. 2025 Oct 7;26(1):342. doi: 10.1186/s13059-025-03773-6. PMID: 41057862; PMCID: PMC12502291.

65. Shinohara M, Kanekiyo T, Yang L, Linthicum D, Shinohara M, Fu Y, Price L, Frisch-Daiello JL, Han X, Fryer JD, Bu G. APOE2 eases cognitive decline during Aging: Clinical and preclinical evaluations. Ann Neurol. 2016 May;79(5):758–774. doi: 10.1002/ana.24628. Epub 2016 Mar 29. PMID: 26933942; PMCID: PMC5010530.

66. Squair JW, Gautier M, Kathe C, Anderson MA, James ND, Hutson TH, Hudelle R, Qaiser T, Matson KJE, Barraud Q, Levine AJ, La Manno G, Skinnider MA, Courtine G. Confronting false discoveries in single-cell differential expression. Nat Commun. 2021 Sep 28;12(1):5692. doi: 10.1038/s41467-021-25960-2. PMID: 34584091; PMCID: PMC8479118.

67. Stoeger T, Grant RA, McQuattie-Pimentel AC, Anekalla KR, Liu SS, Tejedor-Navarro H, Singer BD, Abdala-Valencia H, Schwake M, Tetreault MP, Perlman H, Balch WE, Chandel NS, Ridge KM, Sznajder JI, Morimoto RI, Misharin AV, Budinger GRS, Nunes Amaral LA. Aging is associated with a systemic length-associated transcriptome imbalance. Nat Aging. 2022 Dec;2(12):1191–1206. doi: 10.1038/s43587-022-00317-6. Epub 2022 Dec 9. PMID: 37118543; PMCID: PMC10154227.

68. Stringer C, Wang T, Michaelos M, Pachitariu M. Cellpose: a generalist algorithm for cellular segmentation. Nat Methods. 2021 Jan;18(1):100–106. doi: 10.1038/s41592-020-01018-x. Epub 2020 Dec 14. PMID: 33318659.

69. Sun ED, Zhou OY, Hauptschein M, Rappoport N, Xu L, Navarro Negredo P, Liu L, Rando TA, Zou J, Brunet A. Spatial transcriptomic clocks reveal cell proximity effects in brain ageing. Nature. 2025 Feb;638(8049):160–171. doi: 10.1038/s41586-024-08334-8. Epub 2024 Dec 18. PMID: 39695234; PMCID: PMC11798877.

70. Tabula Muris Consortium. A single-cell transcriptomic atlas characterizes ageing tissues in the mouse. Nature. 2020;583:590–595. doi:10.1038/s41586-020-2496-1.

71. Tung P-Y, Blischak JD, Hsiao CJ, Knowles DA, Burnett JE, Pritchard JK, et al. Batch effects and the effective design of single-cell gene expression studies. Scientific Reports. 2017;7:39921.

72. Uemura E. Age-related changes in neuronal RNA content in rhesus monkeys (Macaca mulatta). Brain Res Bull. 1980 Mar-Apr;5(2):117–9. doi: 10.1016/0361-9230(80)90182-3. PMID: 6155181.

73. Vasconcelos AG, McGuire D, Simon N, Danaher P, Shojaie A. Differential expression analysis for spatially correlated data using smiDE. Genome Biology. 2026;27:21. doi:10.1186/s13059-025-03867-1.

74. Virshup I, Rybakov S, Theis FJ, Angerer P, Wolf FA. AnnData: Annotated data. J Open Source Softw. 2022;7(76):4371. doi: 10.21105/joss.04371.

75. Wang Q, Ding SL, Li Y, Royall J, Feng D, Lesnar P, Graddis N, Naeemi M, Facer B, Ho A, Dolbeare T, Blanchard B, Dee N, Wakeman W, Hirokawa KE, Szafer A, Sunkin SM, Oh SW, Bernard A, Phillips JW, Hawrylycz M, Koch C, Zeng H, Harris JA, Ng L. The Allen Mouse Brain Common Coordinate Framework: A 3D Reference Atlas. Cell. 2020 May 14;181(4):936–953.e20. doi: 10.1016/j.cell.2020.04.007. Epub 2020 May 7. PMID: 32386544; PMCID: PMC8152789.

76. Wang X. Microarray analysis of ageing-related signatures and their expression in tumors based on a computational biology approach. Genomics Proteomics Bioinformatics. 2012 Jun;10(3):136–41. doi: 10.1016/j.gpb.2012.01.001. Epub 2012 Jun 25. PMID: 22917186; PMCID: PMC3586943.

77. Wang Y, Song B, Wang S, Chen M, Xie Y, Xiao G, et al. Sprod for de-noising spatially resolved transcriptomics data based on position and image information. Nature Methods. 2022;19(8):950–958. doi:10.1038/s41592-022-01560-w

78. Wang Y, Zang C, Li Z, Guo CC, Lai D, Wei P. A comparative study of statistical methods for identifying differentially expressed genes in spatial transcriptomics. PLoS Comput Biol. 2026; 22(2): e1013956. 10.1371/journal.pcbi.1013956

79. Wang Z, Zheng Y, Wang F, Zhong J, Zhao T, Xie Q, Zhu T, Ma F, Tang Q, Zhou B, Zhu J. Mfsd2a and Spns2 are essential for sphingosine-1-phosphate transport in the formation and maintenance of the blood-brain barrier. Sci Adv. 2020 May 29;6(22):eaay8627. doi: 10.1126/sciadv.aay8627. PMID: 32523984; PMCID: PMC7259944.

80. Wolf FA, Angerer P, Theis FJ. SCANPY: large-scale single-cell gene expression data analysis. Genome Biol. 2018 Feb 6;19(1):15. doi: 10.1186/s13059-017-1382-0. PMID: 29409532; PMCID: PMC5802054.

81. Wolock SL, Lopez R, Klein AM. Scrublet: Computational Identification of Cell Doublets in Single-Cell Transcriptomic Data. Cell Syst. 2019 Apr 24;8(4):281–291.e9. doi: 10.1016/j.cels.2018.11.005. Epub 2019 Apr 3. PMID: 30954476; PMCID: PMC6625319.

82. Wu C, Macleod I, Su AI. BioGPS and MyGene.info: organizing online, gene-centric information. Nucleic Acids Res. 2013 Jan;41(Database issue):D561–5. doi: 10.1093/nar/gks1114. Epub 2012 Nov 21. PMID: 23175613; PMCID: PMC3531157.

83. Xiang Y, Gu Q, Liu D. Brain Endothelial Cells in Blood-Brain Barrier Regulation and Neurological Therapy. Int J Mol Sci. 2025 Jun 18;26(12):5843. doi: 10.3390/ijms26125843. PMID: 40565303; PMCID: PMC12193000.

84. Ximerakis M, Lipnick SL, Innes BT, Simmons SK, Adiconis X, Dionne D, et al. Single-cell transcriptomic profiling of the aging mouse brain. Nat Neurosci. 2019;22:1696–1708. doi:10.1038/s41593-019-0491-3.

85. Xu C, Lopez R, Mehlman E, Regier J, Jordan MI, Yosef N. Probabilistic harmonization and annotation of single-cell transcriptomics data with deep generative models. Mol Syst Biol. 2021 Jan;17(1):e9620. doi: 10.15252/msb.20209620. PMID: 33491336; PMCID: PMC7829634.

86. Yao Z, van Velthoven CTJ, Kunst M, Zhang M, McMillen D, Lee C, Jung W, Goldy J, Abdelhak A, Aitken M, Baker K, Baker P, Barkan E, Bertagnolli D, Bhandiwad A, Bielstein C, Bishwakarma P, Campos J, Carey D, Casper T, Chakka AB, Chakrabarty R, Chavan S, Chen M, Clark M, Close J, Crichton K, Daniel S, DiValentin P, Dolbeare T, Ellingwood L, Fiabane E, Fliss T, Gee J, Gerstenberger J, Glandon A, Gloe J, Gould J, Gray J, Guilford N, Guzman J, Hirschstein D, Ho W, Hooper M, Huang M, Hupp M, Jin K, Kroll M, Lathia K, Leon A, Li S, Long B, Madigan Z, Malloy J, Malone J, Maltzer Z, Martin N, McCue R, McGinty R, Mei N, Melchor J, Meyerdierks E, Mollenkopf T, Moonsman S, Nguyen TN, Otto S, Pham T, Rimorin C, Ruiz A, Sanchez R, Sawyer L, Shapovalova N, Shepard N, Slaughterbeck C, Sulc J, Tieu M, Torkelson A, Tung H, Valera Cuevas N, Vance S, Wadhwani K, Ward K, Levi B, Farrell C, Young R, Staats B, Wang MM, Thompson CL, Mufti S, Pagan CM, Kruse L, Dee N, Sunkin SM, Esposito L, Hawrylycz MJ, Waters J, Ng L, Smith K, Tasic B, Zhuang X, Zeng H. A high-resolution transcriptomic and spatial atlas of cell types in the whole mouse brain. Nature. 2023 Dec;624(7991):317–332. doi: 10.1038/s41586-023-06812-z. Epub 2023 Dec 13. PMID: 38092916; PMCID: PMC10719114.

87. Zehr KR, Walker MK. Omega-3 polyunsaturated fatty acids improve endothelial function in humans at risk for atherosclerosis: A review. Prostaglandins Other Lipid Mediat. 2018 Jan;134:131–140. doi: 10.1016/j.prostaglandins.2017.07.005. Epub 2017 Aug 9. PMID: 28802571; PMCID: PMC5803420.

88. Zhou T, Xiang L, Liao K, He Y, Zhuang Z, Liu S. stTransfer enables transfer of single-cell annotations to spatial transcriptomics with single-cell resolution. Cell Rep Methods. 2025 Nov 17;5(11):101205. doi: 10.1016/j.crmeth.2025.101205. Epub 2025 Oct 15. PMID: 41101315; PMCID: PMC12664899.

89. Zhou X, Dong K, Zhang S. Integrating spatial transcriptomics data across different conditions, technologies and developmental stages. Nat Comput Sci. 2023 Oct;3(10):894–906. doi: 10.1038/s43588-023-00528-w. Epub 2023 Oct 12. PMID: 38177758.

