## Additional Files for "Scale-Aware Compositional Inference Improves Reproducibility and Uncovers Convergent Aging Programs in Spatial Transcriptomics": Additional_File_2.pdf

### Supplementary Figure 1

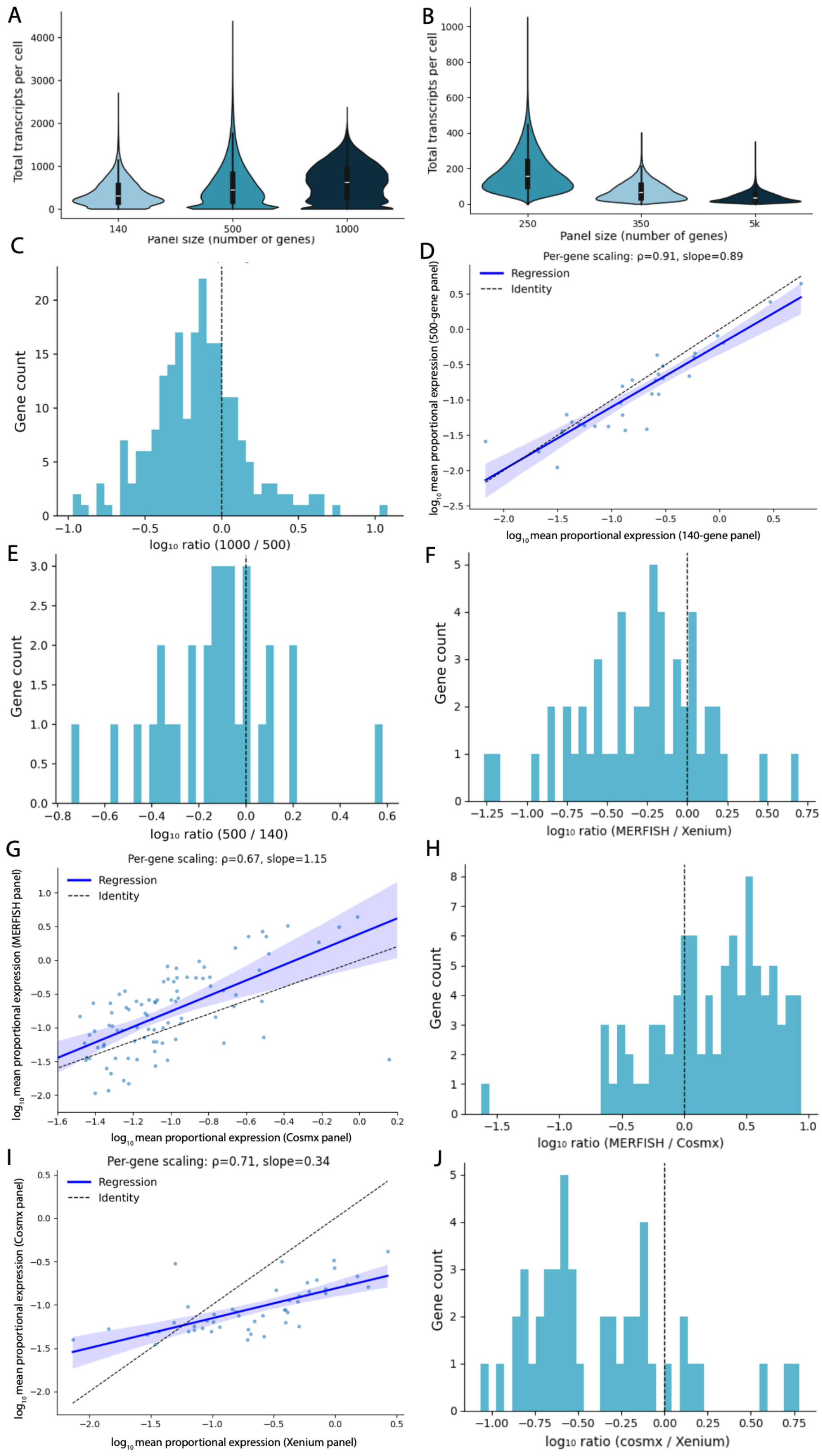

**Figure S1. Additional evidence for scale uncertainty across ST platforms and panel designs.** (a) Total transcripts detected per cell across MERFISH panels of increasing size (140-, 500-, and 1000-gene panels) using all genes in each panel (complementary to Fig. 1A). Transcript abundance increases with panel size but deviates from proportional scaling expectations. (b) Total transcripts detected per cell for genes shared across Xenium panels of different sizes. As observed for MERFISH (Fig. 1A), transcript counts for shared genes vary systematically with panel size despite restricting analysis to the same gene set. (c) Distribution of gene-specific shifts in proportional expression between the 500- and 1000-gene MERFISH panels. Histograms show log10 expression ratios for genes shared between panels, illustrating heterogeneous changes in gene detectability across panel designs. (d) Comparison of $\log_{10}$ mean proportional expression for genes shared between the 140- and 500-gene MERFISH panels. Deviations from the identity line indicate that proportional expression is not preserved across panel compositions. (e) Distribution of gene-specific shifts in proportional expression between the 140- and 500-gene MERFISH panels. Histograms show log10 expression ratios for genes shared between panels. (f) Distribution of gene-specific shifts in proportional expression between MERFISH and Xenium. Histograms show log10 expression ratios for genes shared across platforms, demonstrating heterogeneous platform-dependent differences in gene detectability. (g) Comparison of $\log_{10}$ mean proportional expression for genes shared between MERFISH and CosMx platforms. Deviations from the identity line indicate that proportional expression is not preserved across platforms. (h) Distribution of gene-specifishifts in proportional expression between MERFISH and CosMx. Histograms show log10 expression ratios for genes shared across platforms. (i) Comparison of $\log_{10}$ mean proportional expression for genes shared between Xenium and CosMx platforms. Deviations from the identity line indicate that proportional expression is not preserved across platforms. (j) Distribution of gene-specific shifts in prop between Xenium and CosMx. Histograms show log10 expression ratios for genes shared across platforms.

**Supplementary Figure 2**

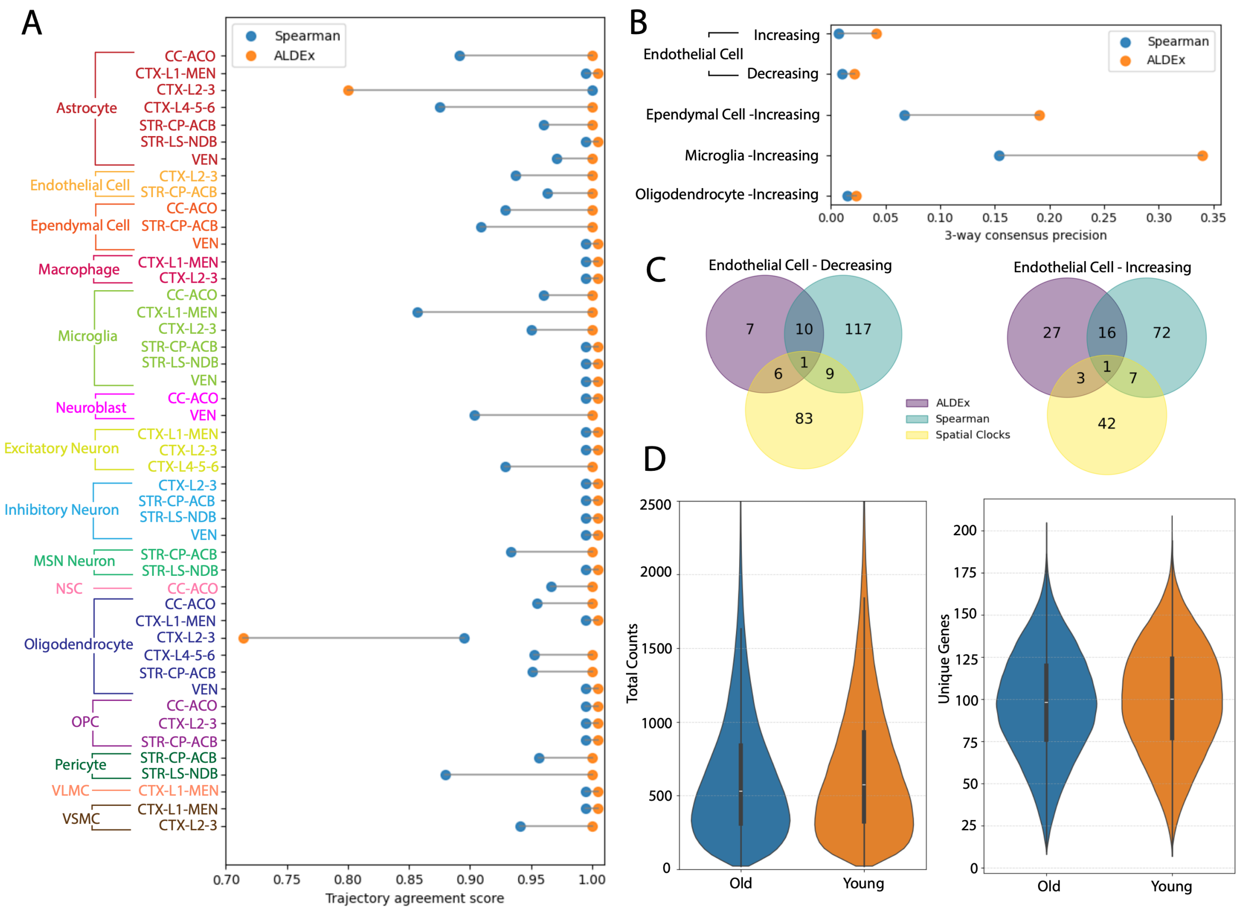


**Figure S2. Supplementary validation analyses in the Sun et al. coronal aging dataset.** (a) Agreement between inferred aging-associated expression changes and published lifespan trajectory classifications from the Sun et al. coronal aging dataset. For each matched cell type × subregion combination, points show the proportion of genes whose inferred direction of change agrees with the corresponding trajectory direction for continuous-age ALDEx and the published Spearman-based analysis. Connected points indicate matched comparisons. (b) Three-way consensus of age-associated GO-BP enrichments across ALDEx, published Spearman-based DE analysis, and spatial ageing clock genes. Connected points indicate matched cell type–direction comparisons, with higher values indicating greater agreement across independent analytical frameworks. (c) Example overlap of enriched GO-BP terms among ALDEx, published Spearman-based DE analysis, and spatial ageing clock genes for decreasing (left) and increasing (right) endothelial-cell ageing programs in the Sun coronal dataset. Although ALDEx identified fewer enriched GO-BP terms overall, a large proportion were shared with both alternative approaches, indicating enrichment for consensus ageing-associated biological programs. (d) Distribution of total transcript counts (left) and detected genes per cell (right) in matched young (3.4–5.4 months) and old (30.9–34.5 months) cohorts from the Sun coronal dataset. Age groups were selected from samples paired within experimental slides to minimize technical confounding. Old cells exhibited modest reductions in both total transcript abundance and the number of detected genes per cell relative to young cells.

#### Supplementary Figure 3

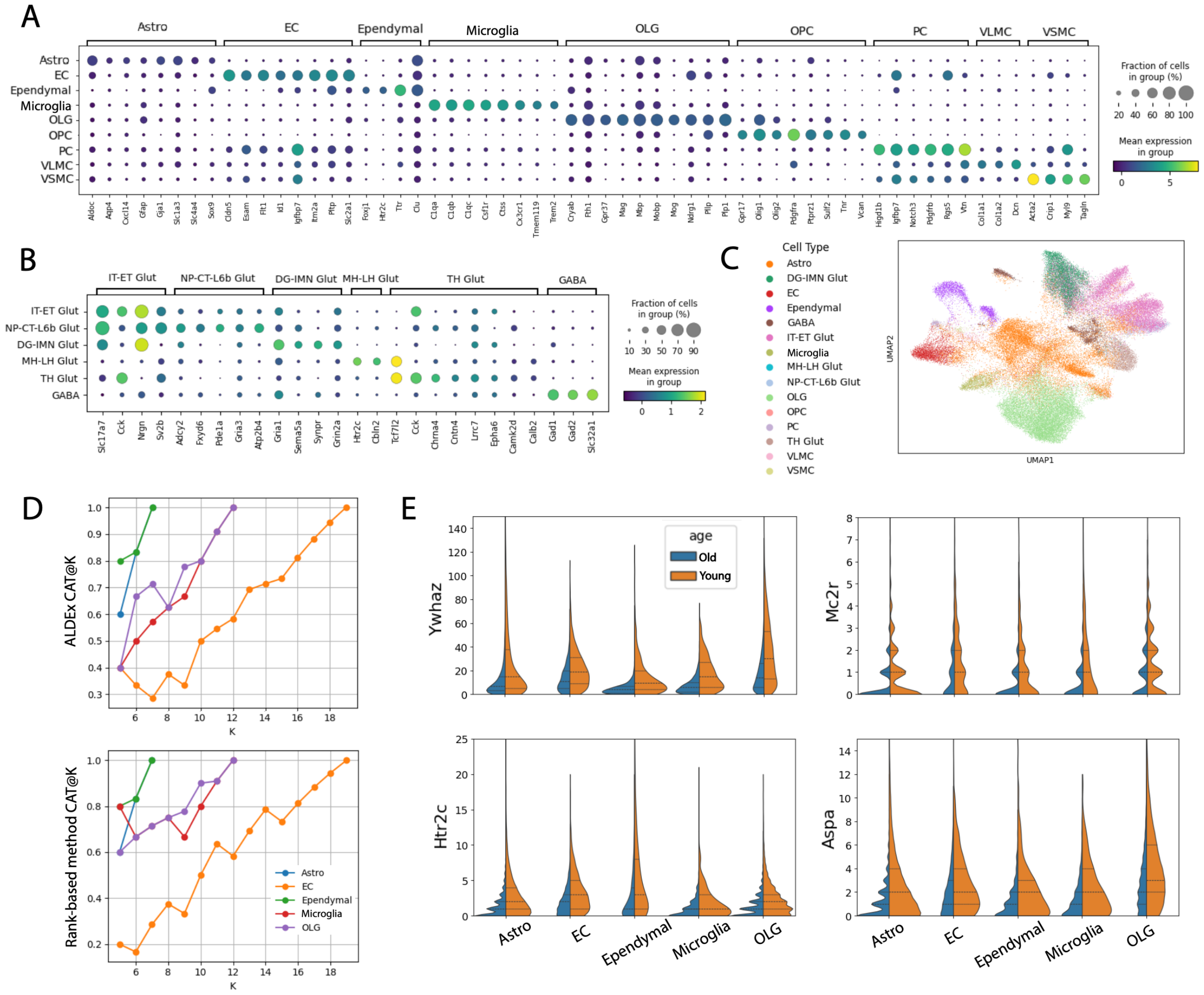


**Figure S3. Cell-type annotation and cross-study reproducibility analyses for the CosMx SenNet aging dataset.** (a) Dot plot of non-neuronal cell-type assignments in the CosMx SenNet aging dataset. Expression of representative marker genes is shown across broad glial, vascular, and immune populations following consensus reference mapping using MapMyCells and scANVI. (b) Dot plot of neuronal cell-class assignments in the CosMx SenNet aging dataset. Representative marker genes are shown for major excitatory and inhibitory neuronal classes following consensus reference mapping. (c) UMAP visualization of CosMx cells colored by final consensus cell-type assignments. Labels were derived using confidence-weighted integration of MapMyCells and scANVI predictions following quality control and filtering.(d) CAT@K analysis comparing overlap among top-ranked aging-associated genes between the primary MERFISH and CosMx datasets across a range of K values. Results are shown separately for ALDEx and the adapted rank-based pseudobulk framework, with higher values indicating greater agreement among highly ranked aging-associated genes across studies. (e) Additional examples of aging-associated genes identified in the independent CosMx SenNet aging dataset. Violin plots show raw transcript abundance distributions in young and aged cells for representative genes exhibiting consistent decreases or increases across multiple non-neuronal populations. Corresponding examples shown in Fig. 4E highlight recurrent age-associated transcriptional changes supported by shifts in observed transcript abundance.

#### Supplementary Table S1: Compositional Analysis Datasets

| Subfigure(s) | Dataset / comparison | Platform | Panel size | Number shared genes used | Source / citation | URL |
|---|---|---|---|---|---|---|
| Fig. 1A; Fig. S1A | MERFISH 140 vs 500 vs 1000 shared-gene panel-size comparison | MERFISH | 140, 500, 1000 | 31 | Internal | Zenodo DOI: 10.5281/zenodo.21420404 |
| Fig. 1B; Fig. S1C | MERFISH 500 vs 1000 gene-level drift | MERFISH | 500, 1000 | 215 | Internal | Zenodo DOI: 10.5281/zenodo.21420404 |
| Fig. S1D–E | MERFISH 140 vs 500 gene-level drift | MERFISH | 140, 500 | 31 | Internal | Zenodo DOI: 10.5281/zenodo.21420404 |
| Fig. 1C; Fig. S1F | MERFISH vs Xenium gene-level drift | MERFISH; Xenium | 500; 250 | 52 | Internal + 10x Genomics Fresh Frozen Mouse Brain Replicates | https://www.10xgenomics.com/datasets/fresh-frozen-mouse-brain-replicates-1-standard |
| Fig. S1B | Xenium panel-size comparison | Xenium | 250, 350, 5k | 150 | 10x Genomics public datasets:<br>1) Fresh Frozen Mouse Brain Replicates<br>2) Xenium In Situ Analysis of Alzheimer's Disease Mouse Model Brain Coronal Sections from One Hemisphere Over a Time Course<br>3) Fresh Frozen Mouse Brain Hemisphere with 5K Mouse Pan Tissue and Pathways Panel | 1) https://www.10xgenomics.com/datasets/fresh-frozen-mouse-brain-replicates-1-standard<br>2) https://www.10xgenomics.com/datasets/xenium-in-situ-analysis-of-alzheimers-disease-mouse-model-brain-coronal-sections-from-one-hemisphere-over-a-time-course-1-standard<br>3) https://www.10xgenomics.com/datasets/xenium-prime-fresh-frozen-mouse-brain |
| Fig. S1G–H | MERFISH vs CosMx gene-level drift | MERFISH; CosMx | 500; 1000 | 92 | Internal + SenNet SNT638.MWRV.378 | doi:10.60586/SNT638.MWRV.378 |
| Fig. S1I–J | Xenium vs CosMx gene-level drift | Xenium; CosMx | 250; 1000 | 47 | 10x Genomics Fresh Frozen Mouse Brain Replicates<br>SenNet SNT638.MWRV.378 | https://www.10xgenomics.com/datasets/fresh-frozen-mouse-brain-replicates-1-standard<br>doi:10.60586/SNT638.MWRV.378 |
| Fig. 1E–G | Matched MERFISH coronal sections | MERFISH | 500 | 487 | Internal | Zenodo DOI: 10.5281/zenodo.21420404 |
| Fig. 1D | Aging associated gene sets from 3 mouse brain single-cell transcriptomic studies | 1) Tabula Muris Senis: 10x Genomics Chromium; Smart-seq2<br>2) Broad Institute: 10x Genomics Chromium 3′ v2<br>3) Allen Institute: 10x Genomics Chromium 3′ v3 | Whole transcriptome | 88 (oligodendrocytes), 24 (endothelial cells), 33 (microglia) | 1) Tabula Muris Senis: Tabula Muris Consortium (2020), processed gene-count and metadata record<br>2) Broad Institute: Ximerakis et al. (2019), Supplementary Data: “Differential gene expression data between young and old cell types”<br>3) Allen Institute: Jin et al. (2025), Supplementary Data workbook: “supertype table” sheet | 1) Tabula Muris Senis: https://doi.org/10.1038/s41586-020-2496-1 processed data record: https://doi.org/10.6084/m9.figshare.8273102.v2<br>2) Broad Institute: https://doi.org/10.1038/s41593-019-0491-3<br>3) Allen Institute: https://doi.org/10.1038/s41586-024-08350-8 |

Table S2: Sun et al. effect-size concordance metrics

| Cell type | N genes | ALDEx Pearson r | ALDEx Spearman ρ | Pseudobulk Person r | Pseudobulk Spearman ρ |
|---|---|---|---|---|---|
| Isocortex Astro | 22 | 0.56 | 0.64 | 0.51 | 0.48 |
| Isocortex Ec | 26 | 0.39 | 0.4 | 0.27 | 0.26 |
| Isocortex Immune | 24 | 0.38 | 0.36 | 0.44 | 0.51 |
| Isocortex Olg | 25 | 0.52 | 0.43 | 0.64 | 0.52 |
| Isocortex Opc | 26 | 0.21 | 0.4 | 0.31 | 0.33 |
| Isocortex Pc | 26 | 0.36 | 0.42 | 0.3 | 0.46 |
| Isocortex Vsmc | 24 | -0.25 | -0.45 | -0.36 | -0.35 |

Table S3: SenNet Cosmx dataset effect-size concordance metrics

| Cell type | N genes | ALDEx Pearson r | ALDEx Spearman ρ | Pseudobulk Pearson r | Pseudobulk Spearman ρ |
|---|---|---|---|---|---|
| Astrocyte | 7 | -0.61 | -0.43 | -0.11 | -0.07 |
| Endothelial Cell | 19 | -0.03 | -0.07 | -0.07 | -0.12 |
| Ependymal | 7 | 0.69 | 0.71 | -0.44 | -0.43 |
| Immune | 12 | 0.05 | 0.08 | 0.45 | 0.25 |
| Oligodendrocyte | 12 | -0.24 | -0.01 | -0.35 | 0.1 |
